# Dual Engram Architecture within a Single Striatal Cell Type Distinctly Controls Alcohol Relapse and Extinction

**DOI:** 10.64898/2026.01.13.699375

**Authors:** Xueyi Xie, Yufei Huang, Ruifeng Chen, Zhenbo Huang, Himanshu Gangal, Jiayi Lu, Adelis M. Cruz, Anita Chaiprasert, Emily Yu, Niko Hernandez, Valerie Vierkant, Xuehua Wang, Rachel J. Smith, Jun Wang

**Affiliations:** Department of Neuroscience and Experimental Therapeutics, College of Medicine, Texas A&M University Health Science Center; Bryan, Texas 77807, United States; Institute for Neuroscience, Texas A&M University; College Station, Texas 77843, United States; Department of Psychological and Brain Sciences, Texas A&M University; College Station, Texas 77843, United States

**Keywords:** engram ensemble, relapse, extinction, alcohol use disorder, direct-pathway medium spiny neurons (dMSNs), dorsomedial striatum (DMS), operant self-administration (OSA), ArcTRAP/FosTRAP, long-term potentiation (LTP), striosome, mu-opioid receptor (MOR)

## Abstract

Relapse is a major obstacle in the treatment of alcohol and drug addiction and is thought to be driven by persistent drug-associated memories formed during their use. Behavioral therapies such as extinction training can reduce relapse and are proposed to work by creating a competing memory trace. However, where and how these opposing memories are stored in the brain is unknown. Here, we show that two anatomically and functionally distinct engram ensembles within the same genetically defined striatal cell type, direct-pathway medium spiny neurons (dMSNs), encode these opposing memories. Using engram-tagging tools in mice, we found that the acquisition of operant alcohol learning recruits a dMSN ensemble enriched in the striatal matrix compartment that stores alcohol-associated memories and whose activation selectively promotes relapse. Conversely, extinction of alcohol seeking recruits a dMSN ensemble enriched in the striosome compartment that stores extinction-related memories and whose activation suppresses relapse. Furthermore, we reveal that the physical memory trace storing the relapse-promoting memory is embedded within persistently strengthened corticostriatal synapses engaged during learning, and that artificially reproducing this plasticity is sufficient to trigger relapse-like behavior. These findings uncover a dual-engram architecture within dMSNs that governs relapse and extinction, providing a mechanistic framework for understanding how competing memories regulate drug-seeking behavior.

**Highlights:**

1. Acquisition and extinction of alcohol learning recruits distinct dMSN ensembles.
2. Acquisition recruits matrix-enriched dMSN ensembles to promote relapse.
3. Extinction recruits striosome-enriched dMSN ensembles to suppress relapse.
4. Engram dMSNs show lasting synaptic potentiation and mimicking this potentiation triggers relapse.

## Introduction

Drug and alcohol addiction arises when normal learning processes are hijacked by addictive substances, resulting in the formation of maladaptive drug-related memories^1–3^. These drug-related memories are enduring and lead to recurrent episodes of relapse, presenting a significant challenge in addiction treatment^2^. Behavioral therapies such as extinction training can reduce relapse by forming a competing memory that suppresses drug seeking, but their clinical impact remains limited^1,4–9^, partly because the underlying cellular substrates of these opposing memories are poorly understood.

Memories are thought to be encoded by engram ensembles, which are specific groups of neurons that undergo learning-induced synaptic changes and are reactivated during memory retrieval^10–13^. Most engram studies have focused on Pavlovian fear learning, identifying engram ensembles in the hippocampus, cortex, and amygdala that store and retrieve fear memories^10–12,14^. In contrast, less is known about the engram ensembles that encode persistent instrumental memories for addictive substances^15,16^, despite the fact that human relapse often results from voluntary, goal-directed behaviors as much as it does from Pavlovian cues^17,18^.

The dorsomedial striatum (DMS) is essential for both instrumental learning and relapse to drug and alcohol seeking^19–22^. Within the DMS, direct-pathway medium spiny neurons (dMSNs) promote reinforcement^23–26^, are preferentially activated by addictive substances^27–30^, and are required for relapse^31–34^. Traditionally viewed as a uniform “Go” population, dMSNs are organized into neurochemically distinct matrix and striosome compartments with different connectivity and behavioral roles: matrix dMSNs are associated with reward and positive reinforcement, whereas striosomal dMSNs are linked to negative reinforcement and behavioral inhibition^35–38^. This raises the possibility that relapse- and extinction-related memories might arise from distinct compartment-specific dMSN ensembles yet remain within the same genetically defined cell type—a concept largely overlooked by striatal circuit models.

Here, we test this idea using an operant alcohol self-administration model combined with activity-dependent ensemble tagging, circuit manipulations, and synaptic physiology. We identify a dual-engram architecture within DMS dMSNs, in which matrix-enriched ensembles encode relapse-promoting memories, striosome-enriched ensembles encode extinction memories, and the physical substrate of the relapse engram resides in specific corticostriatal synapses. This reveals how anatomically distinct ensembles within a single cell type can exert opposing control over complex, learned behaviors.

## Results

### Acquisition and extinction of alcohol-associated memories recruit distinct striatal dMSN ensembles

We first asked whether acquisition and extinction of operant alcohol seeking recruit distinct ensembles of DMS dMSNs. To test this, we employed ArcTRAP mice, which enables activity-dependent neuronal tagging^39^, and the operant alcohol self-administration paradigm, which produces robust alcohol-associated memories and models key features of human relapse behavior^18^. It is well established that retrieval of associative memories selectively reactivates neuronal ensembles originally activated during the corresponding learning episode^12,14^. This reactivation is memory-specific: ensembles recruited by one type of learning are less likely to be reactivated during retrieval of a different memory^40–42^. Thus, “reactivation” serves as a key method to determine whether the same group of neurons is involved in different memory traces. Based on this principle, we hypothesized that dMSNs activated during the acquisition of operant alcohol learning would be preferentially reactivated during alcohol memory retrieval, whereas those activated during extinction would not (Fig. 1A).

**Figure 1.**
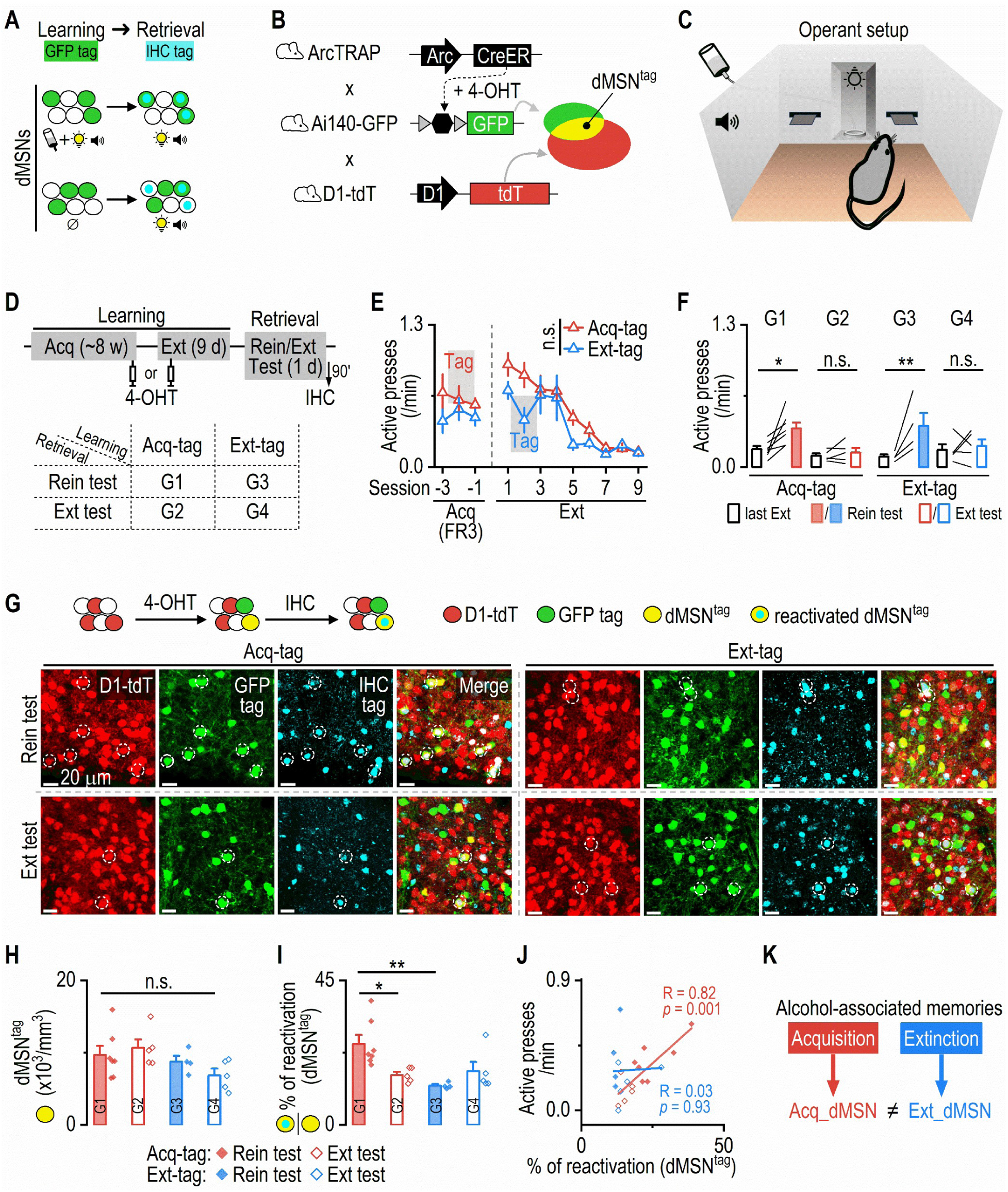
Acquisition- and extinction-recruited dMSNs are differentially reactivated during cue-induced reinstatement of alcohol seeking. ***A***, Schematic illustrating the hypothesis that neurons activated during acquisition of alcohol-associated learning (tagged with GFP) are more likely to be reactivated during retrieval of the alcohol-cue memory (detected via immunohistochemistry [IHC]), compared to neurons activated during extinction learning, which are less likely to be reactivated during alcohol-cue retrieval. ***B***, Mechanism of activity-dependent GFP expression in tagged neurons and tdTomato (tdT) expression in dMSNs of ArcTRAP;Ai140-GFP;D1-tdT mice. Neurons co-expressing Ai140-GFP and D1-tdT are designated as dMSN^tag^. ***C***, Schematic of the operant alcohol self-administration paradigm. Two retractable levers flank a central magazine with an alcohol delivery port and a cue light above. A tone generator is positioned on the chamber’s back wall. ***D***, Top: ArcTRAP;Ai140-GFP;D1-tdT mice received ∼8 weeks of operant alcohol self-administration (acquisition), followed by 9 days of extinction training. Neurons were tagged with 4-OHT either on the second-to-last day of acquisition (Acq-tag) or on the second day of extinction (Ext-tag). After completing extinction training, mice were either tested for cue-induced reinstatement (Rein) or for an additional extinction session (Ext). Ninety minutes after the test, brains were collected for c-Fos IHC. Group assignments: Group 1 (G1): Acq-tag to Rein; Group 2 (G2): Acq-tag to Ext; Group 3 (G3): Ext-tag to Rein; Group 4 (G4): Ext-tag to Ext. ***E***, Acq-tag and Ext-tag groups exhibited similar active lever presses during acquisition and extinction phases (two-way RM ANOVA: Group × Session, *F*_(11,209)_ = 1.27, *p* = 0.24). n.s., not significant. ***F***, Groups tested for reinstatement (G1, G3) exhibited increased active presses, while extinction-tested groups (G2, G4) did not. Two-way RM ANOVA: Group × Test, *F*_(3,17)_ = 3.27, *p* = 0.047; Sidak’s multiple comparisons test: last Ext vs. Test for G1: *t* = 3.40, \**p* = 0.01; G2: *t* = 0.43, *p* = 0.98; G3: *t* = 3.82, \*\**p* = 0.006; G4: *t* = 0.54, p = 0.97. ***G***, Representative confocal images showing tagged dMSNs reactivated across the four groups. Circled neurons represent tagged dMSNs that were reactivated during the test. ***H***, Acq-tag and Ext-tag groups had similar densities of dMSN^tag^. Two-way ANOVA: Group, *F*_(1,17)_ = 1.49, *p* = 0.06. ***I***, The Acq-tag group tested for reinstatement exhibited the highest reactivation rate of dMSN^tag^. Two-way ANOVA: Group × Test, *F*_(1,17)_ = 8.38, *p* = 0.01; Sidak’s multiple comparisons test: G1 vs G2: *t* = 3.00, \**p* = 0.047; G1 vs G3: *t* = 3.76, \*\**p* = 0.009. *J*, Reactivation of Acq-tag dMSN^tag^ was positively correlated with active presses during the test, whereas Ext-tag dMSN^tag^ reactivation was not. Liner regression. ***K***, Summary diagram illustrating that distinct dMSN populations are recruited by acquisition versus extinction of alcohol-associated memories. dMSNs tagged during acquisition (Acq_dMSN, red) are preferentially reactivated during alcohol-cue memory retrieval, whereas dMSNs tagged during extinction (Ext_dMSN, blue) are less reactivated during the same retrieval. n = 7 mice (Acq-tag, Rein test), 5 mice (Acq-tag, Ext test), 4 mice (Ext-tag, Rein test), and 5 mice (Ext-tag, Ext test).

To begin, we generated and trained triple transgenic mice (ArcTRAP;Ai140-GFP;D1-tdT) to lever-press alcohol (20% v/v) in operant chambers for ∼8 weeks (Fig. 1B-1D). These mice allow us to label learning “tagged” neurons with GFP, dMSNs with tdTomato (tdT), and tagged dMSNs (dMSN^tag^) with the overlap of both markers (Fig. 1B). The training involved a fixed ratio (FR) schedule that eventually required three active presses (FR3) to trigger an alcohol delivery, which was accompanied by discrete cues (light+tone). Mice then underwent 9 d of extinction training, in which active presses no longer resulted in alcohol or cue delivery^43,44^. Neurons active during either acquisition or extinction were tagged with GFP by injecting 4-hydroxytamoxifen (4-OHT, intraperitoneal, 50 mg/kg) at specific time points. One group received 4-OHT during an FR3 session (acquisition-tagged; Acq-tag), tagging neurons activated during acquisition of operant alcohol learning. Another group received 4-OHT during the second day of extinction training (extinction-tagged; Ext-tag), tagging neurons activated during extinction (Fig. 1D, 1E). Both groups showed comparable acquisition and extinction performance (Fig. 1E). To assess memory-specific ensemble reactivation, mice underwent either a cue-induced reinstatement test (retrieval of the alcohol memory) or an extinction test (retrieval of the extinction memory) (Fig. 1D). The reinstatement, but not extinction, test induced robust increases in active presses in both groups (G1 and G3; Fig. 1F), indicating successful retrieval of alcohol memories during reinstatement. Ninety minutes after the test, mice were sacrificed for c-Fos immunohistochemistry to label neurons activated during reinstatement or extinction testing, and to assess the reactivation of learning-recruited dMSNs (dMSN^tag^) by examining its overlap with c-Fos (Fig. 1D, 1G).

Interestingly, both groups showed similar densities of tagged dMSNs (Fig. 1H; Fig. S1A, S1B), suggesting that both acquisition and extinction of operant alcohol learning recruited dMSNs. Crucially, acquisition-recruited dMSNs (GFP^+^ & tdT^+^) exhibited a higher reactivation rate [(c-Fos^+^ & GFP^+^ & tdT^+^) / (GFP^+^ & tdT^+^)] during reinstatement than during extinction (G1 vs. G2; Fig. 1I; Fig. S1D, S1E). This suggests that acquisition-recruited dMSNs were more likely to reactivate during alcohol memory retrieval than during extinction memory retrieval. Conversely, extinction-recruited dMSNs exhibited a lower reactivation rate than acquisition-recruited dMSNs during reinstatement (G1 vs. G3; Fig. 1I; Fig. S1D, S1E). This indicates that dMSNs recruited by extinction did not participate in alcohol memory retrieval. Strikingly, reactivation of acquisition-tagged dMSNs strongly correlated with their alcohol-seeking behavior during the tests (Fig. 1J), whereas no such correlation was observed in extinction-tagged mice.

Together, these findings demonstrate that both the acquisition and extinction of alcohol-associated memories recruit ensembles within the same genetically defined striatal cell type, dMSNs, yet through distinct, experience-defined populations (Fig. 1K). Additionally, their divergent reactivation patterns during alcohol-memory retrieval indicates that they represent separate neuronal ensembles within the DMS.

### Acquisition- and extinction-recruited dMSNs differ in anatomical distribution, circuit output, and behavioral function

Having shown that acquisition and extinction of operant alcohol learning recruit distinct populations of DMS dMSNs, we next asked whether these ensembles could be distinguished by their spatial distribution, projection patterns, and behavioral roles. While dMSNs are traditionally viewed as “Go” neurons that promote actions^45^, they are not a uniform population. Instead, they are organized into two neurochemically and functionally distinct compartments: matrix and striosomes^35^. Notably, striosomal dMSNs express mu-opioid receptors (MORs)^46,47^, directly inhibit dopamine neurons in the substantia nigra pars compacta (SNc), and discourage actions^27,36–38,47–49^. In contrast, matrix dMSNs, which represent the canonical dMSN population, project to the substantia nigra pars reticulata (SNr) and promote behavioral output upon activation^36,38^. Based on these distinctions, we hypothesized that extinction-recruited dMSNs may be enriched in the striosomal compartment and exert a stronger inhibitory effect on dopamine neurons than acquisition-recruited dMSNs (Fig. 2A).

**Figure 2.**
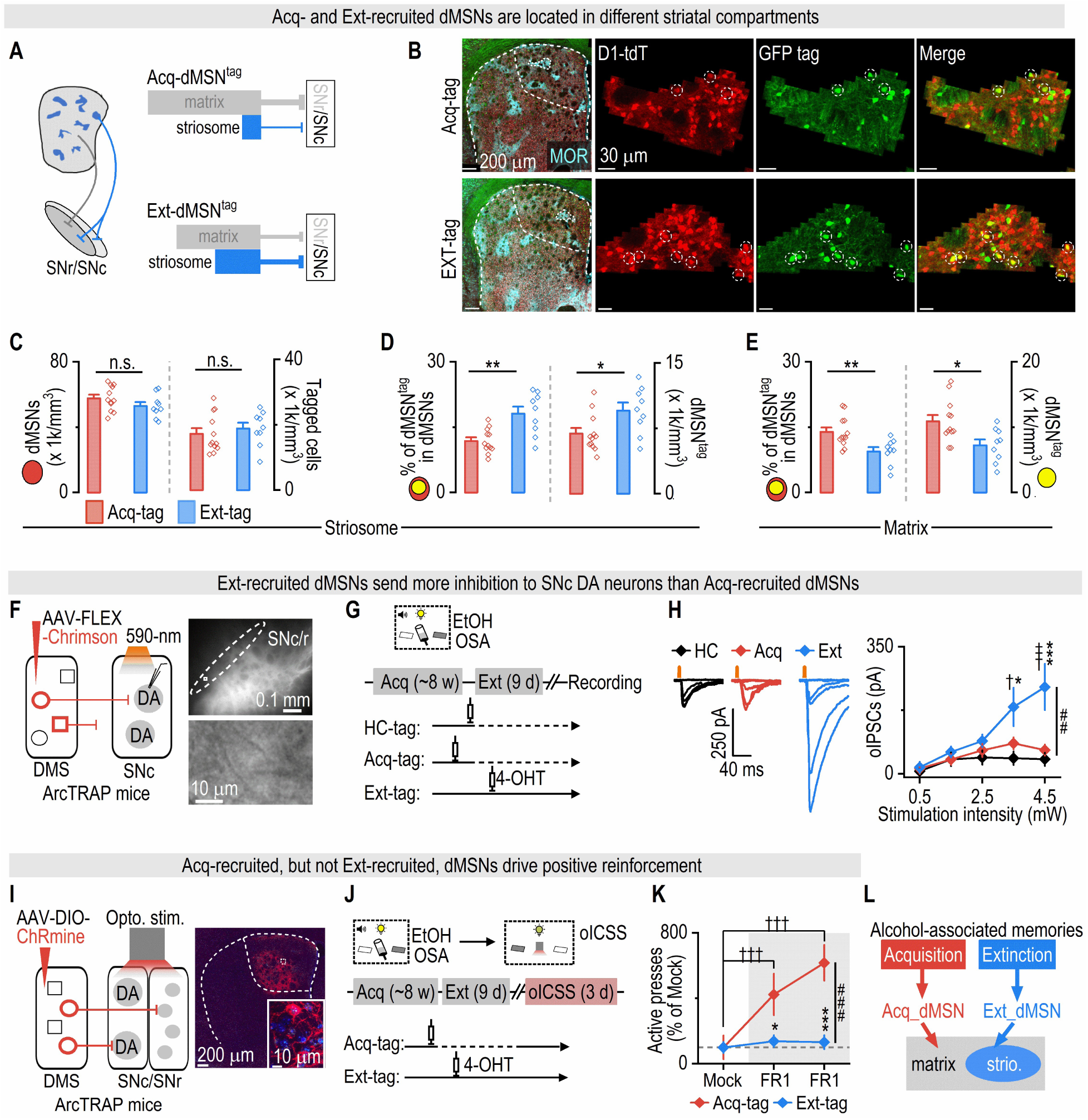
Acquisition- and extinction-recruited dMSNs differ in anatomical distribution, circuit output, and behavioral function. ***A***, Schematic illustration of the hypothesis: extinction recruits a distinct striatal population from acquisition, containing more striosomal dMSNs that send direct inhibitory inputs to SNc dopaminergic neurons. ***B***, Sample confocal images showing dMSN^tag^ in MOR-expressing striosomes in Acq-tag versus Ext-tag mice. Sections were from the same mice in Figure 1 and stained with anti-MOR antibody. ***C***, Both Acq-tag and Ext-tag groups showed similar densities of dMSNs (left) and tagged neurons (right) in the DMS striosome compartment. Left: unpaired t test: *t*_(19)_ = 1.47, *p* = 0.16. Right: unpaired t test: *t*_(19)_ = 0.66, *p* = 0.52. ***D***, The Ext-tag group had a higher percentage of striosomal dMSNs being recruited (left; unpaired t test: *t*_(19)_ = 3.76, \*\**p* = 0.001) and more striosomal dMSN^tag^ density (right; unpaired t test: *t*_(19)_ = 2.51, \**p* = 0.02) than did Acq-tag group. ***E***, The Acq-tag group had a higher percentage of matrix dMSNs being recruited (left; unpaired t test: *t*_(19)_ = 3.11, \*\**p* = 0.006) and more matrix dMSN^tag^ density (right; unpaired t test: *t*_(19)_ = 2.74, \**p* = 0.013) than did Ext-tag group. ***F***, Left: ArcTRAP mice were infused with AAV-FLEX-Chrimson-tdT into the DMS. Optically induced inhibitory postsynaptic currents (oIPSCs) were recorded from SNc DA neurons under 590-nm light stimulation. In the DMS, circle represents dMSN and square represents non-dMSN. Right: Sample image showing tdT+ fibers in the SNc/SNr region, with an enlarged view depicting a recorded DA neuron in the ventromedial SNc. ***G***, Schematic of the experiment timeline. ArcTRAP mice were trained for EtOH OSA for ∼8 weeks. One group received 4-OHT in the home-cage during two consecutive days, 24 h after the last acquisition session (HC-tag). Another group received 4-OHT during the last two acquisition sessions (Acq-tag). Mice in the HC-tag and Acq-tag groups then returned to chambers for 30 min daily, without lever access, for 9 d. A final group underwent 9-d extinction training, receiving 4-OHT during the 2nd and 3rd extinction sessions (Ext-tag). Electrophysiological recordings were conducted two weeks after tagging. ***H***, SNc DA neurons exhibited the highest sIPSC frequencies in the Ext-tag group (Kruskal-Wallis test: *H*_(2)_ = 13.94, *p* < 0.001; Dunn’s multiple comparisons test: HC-tag vs. Ext-tag: *z* = 3.01, \*\**p* = 0.0078; Acq-tag vs. Ext-tag: *z* = 3.38, \*\**p* = 0.0022). ***I***, SNc DA neurons in the Ext-tag group received the strongest oIPSCs from dMSN^tag^ compared to those in the HC-tag and Acq-tag groups. Two-way RM ANOVA: Group × Intensity: *F*_(8,136)_ = 2.84, ^##^*p* = 0.006; Tukey’s multiple comparisons test: HC-tag vs. Ext-tag: q = 3.91, ^†^*p* = 0.017; q = 5.51, ^†††^*p* = 0.0004 under the same stimulation intensity; Acq-tag vs. Ext-tag: q= 3.37, \**p* = 0.046; q = 5.85, \*\*\**p* = 0.0002 under the same stimulation intensity. ***J***, ArcTRAP mice were infused with AAV-DIO-ChRmine-oScarlet into the DMS, and optical fibers were implanted above the SNc/SNr region to stimulate terminals of dMSN^tag^. In the DMS, circle represents dMSN and square represents non-dMSN. ***K***, Schematic of the experiment timeline. Mice were trained for EtOH OSA for ∼8 weeks. One group received 4-OHT during the last two acquisition sessions (Acq-tag), while another group received 4-OHT during the 2nd and 3rd extinction sessions (Ext-tag). Mice in the Acq-tag group remained in the chambers for 30 min daily without lever access for 9 d. Two weeks after extinction, all mice were returned to the chambers for optical intracranial self-stimulation (oICSS) with red light, which included 1 d of mock induction and 2 d of FR1. During oICSS, the previously inactive lever for alcohol became the active lever for light stimulation, and vice versa for the previous active lever. During the mock session, all task parameters (cue light, time-out, lever configuration) were identical to the FR1 sessions, except that active presses did not trigger optical stimulation, allowing measurement of baseline responding before training. ***L***, Mice in the Acq-tag group gradually increased active presses for light stimulation, whereas mice in the Ext-tag group did not. Active presses (normalized to the mock session) were higher in the Acq-tag group than the Ext-tag group. Two-way RM ANOVA: Group × Session: *F*_(2,14)_ = 20.31, ^###^*p* < 0.001. Sidak’s multiple comparisons test: Acq-tag vs. Ext-tag in 1^st^ FR1: *t* = 2.84, \**p* = 0.029; Acq-tag vs. Ext-tag in 2^nd^ FR1: *t* = 4.83, \*\*\**p* < 0.001; mock vs. 1^st^ FR1 in Acq-tag: *t* = 5.67, ^†††^*p* < 0.001; mock vs. 2^nd^ FR1 in Acq-tag: *t* = 9.06, ^†††^*p* < 0.001. ***M***, Summary diagram illustrating that acquisition-recruited dMSNs are predominantly located in the matrix compartment, provide minimal inhibitory input to SNc dopamine neurons, and drive positive reinforcement. In contrast, extinction-recruited dMSNs are enriched in the striosome compartment, send strong inhibitory input to SNc dopamine neurons, and do not support reinforcement behavior. n = 12 mice (Acq-tag; C-E), 9 mice (Ext-tag; C-E), 7 neurons from 3 mice (7/3; HC-tag; H), 13/5 (Acq-tag; H), 17/5 (Ext-tag; H), 4 mice (Acq-tag; K), and 5 mice (Ext-tag; K).

First, we compared the anatomical distribution of acquisition- and extinction-recruited dMSNs across the striosome and matrix compartments. Striosomes were identified by MOR immunostaining, and dMSN^tag^ distribution was examined in triple transgenic mice (ArcTRAP;Ai140-GFP;D1-tdT) tagged during either acquisition or extinction (Fig. 2B; same mice as in Fig. 1D). The two groups showed similar overall densities of total dMSNs and total tagged cells within striosomes (Fig. 2C). However, extinction recruited a greater percentage of striosomal dMSNs, resulting in a higher density of dMSN^tag^ in extinction-tagged mice compared with acquisition-tagged mice (Fig. 2D). Conversely, in the matrix compartment, acquisition recruited a greater percentage of matrix dMSNs, leading to a higher density of dMSN^tag^ in the acquisition group than in the extinction group (Fig. 2E). Together, these findings suggested that two ensembles differ in their anatomical localization: acquisition preferentially recruited matrix-enriched dMSNs, whereas extinction recruited a striosome-enriched dMSN population.

Given that striosomal dMSNs directly inhibit SNc dopamine neurons, whereas matrix dMSNs do not, we next asked whether extinction-recruited dMSNs exerted greater inhibitory control over dopamine neurons than acquisition-recruited dMSNs. To selectively measure dMSN^tag^-to-dopaminergic transmission, we expressed Chrimson in DMS tagged neurons using ArcTRAP mice and recorded inhibitory postsynaptic currents (IPSCs) in ventromedial SNc dopaminergic neurons (Fig. 2F)^27^. Mice were trained for operant alcohol learning until they reached stable FR3 performance (Fig. S2A) and were then divided into three groups for tagging: in the home-cage 24 h after the last acquisition session (HC-tag), during the last two sessions of acquisition (Acq-tag), or during the second and third sessions of extinction training (Ext-tag) (Fig. 2G). Only the Ext-tag group underwent 9 d of lever-extinction training (Fig. S2B), while the HC-tag and Acq-tag groups returned to the chamber daily (for 9 d) without lever extinction. Spontaneous IPSCs in dopaminergic neurons had similar amplitudes but significantly higher frequencies in the Ext-tag group (Fig. 2H; Fig. S2C), indicating increased inhibitory input to dopaminergic neurons after lever-extinction training. We then selectively measured optically induced IPSCs from dMSN^tag^ by delivering 590-nm stimulation. A higher proportion of dopaminergic neurons in the Ext-tag group exhibited optically induced IPSCs, as compared to the other two groups (Fig. S2D), with the largest amplitudes observed in the Ext-tag group (Fig. 2I). These results indicate that extinction-recruited dMSNs, which were enriched in the striosomal compartment, produced stronger inhibition of dopaminergic neurons than did acquisition-recruited dMSNs—consistent with the known direct connectivity between striosomal dMSNs and SNc dopamine neurons^35^.

Given that extinction-recruited dMSNs strongly inhibited dopaminergic neurons, and striosomal dMSNs are associated with behavioral inhibition^37^, we next tested whether optical self-stimulation of acquisition- versus extinction-recruited dMSNs would differentially drive operant reinforcement. We expressed ChRmine in either acquisition-or extinction-tagged striatal neurons in ArcTRAP mice and implanted fibers over the SNc/SNr, allowing optical stimulation of dMSN^tag^ terminals in the direct-pathway output area (Fig. 2J). Mice were trained for operant alcohol learning followed by extinction, with tagging during either the last two acquisition sessions (Acq-tag) or the second/third extinction sessions (Ext-tag) (Fig. 2K; Fig. S2E). Two weeks later, mice returned to the chamber for optogenetic intracranial self-stimulation (oICSS) using 620-nm light stimulation. The previous inactive lever for alcohol now became the active lever for oICSS, and vice versa (Fig. 2K). Notably, the Acq-tag group progressively increased active presses over 3 d, whereas the Ext-tag group did not (Fig. 2L). These results demonstrate that acquisition-recruited dMSNs, which are enriched in the matrix, supported operant reinforcement, whereas extinction-recruited dMSNs, which are enriched in the striosome, did not.

Together, these findings demonstrate that extinction training recruits a striosome-enriched dMSN population that is anatomically distinct from acquisition-recruited matrix dMSNs, sends stronger output to dopamine neurons, and fails to support operant reinforcement, suggesting that these two ensembles encode opposing alcohol-associated memories (Fig. 2M).

### Alcohol acquisition-recruited dMSNs are required for alcohol, but not sucrose, relapse

Having demonstrated that acquisition- and extinction-recruited dMSNs constitute anatomically and functionally distinct populations within the DMS, we next asked whether these ensembles serve as memory-specific engrams, differentially controlling retrieval of alcohol-associated memories. Given that acquisition-recruited dMSNs were preferentially reactivated during alcohol memory retrieval and correlated with relapse, we hypothesized that they constitute a dedicated engram required for the retrieval of cue-alcohol memories driving relapse. Conversely, extinction-recruited dMSNs, which preferentially occupy the striosomal compartment and inhibit dopamine neurons, may represent a separate engram promoting extinction memory retrieval and suppressing relapse. To test this, we selectively inhibited acquisition-recruited dMSNs during cue-induced reinstatement and assessed their causal role in driving alcohol relapse (Fig. 3A).

**Figure 3.**
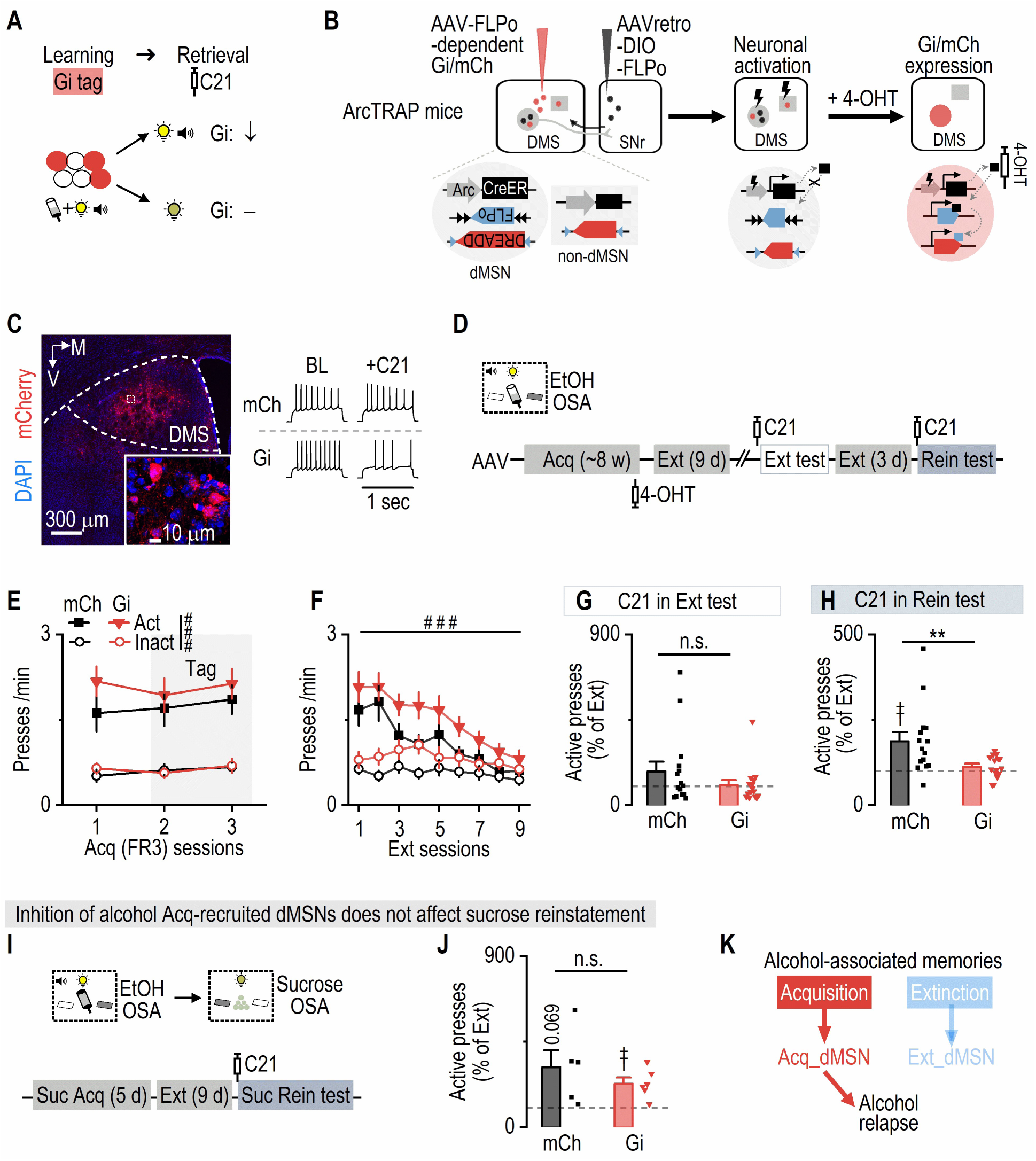
Chemogenetic inhibition of alcohol acquisition-recruited dMSNs suppresses cue-induced reinstatement of alcohol-, but not sucrose-, seeking. ***A*,** Schematic illustrating the hypothesis: chemogenetic inhibition of alcohol acquisition-tagged neurons (expressing Gi) suppresses the retrieval of the same cue-alcohol memory (top), while leaving other memories unaffected (bottom). ***B*,** Schematic of dual-virus infusions in ArcTRAP mice, enabling targeted DREADD expression in acquisition-recruited dMSNs. The SNr was infused with AAVretro-DIO-FLPo, which is retrogradely transported to the DMS only in dMSNs. The DMS was then infused with FLPo-dependent DREADDs (AAV-fDIO-hM4Di-mCherry [Gi] or AAV-fDIO-mCherry [mCh]). ***C*,** Representative confocal images (left) show hM4Di-mCherry^+^ neurons in the DMS. Electrophysiological recordings (right) demonstrate that C21 bath application did not alter evoked firing rates in mCh-tagged neurons but reduced evoked firing rates in Gi-tagged neurons under identical current injections. M, medial; V, ventral. ***D*,** Experimental timeline. Following virus infusion, ArcTRAP mice underwent ∼8 weeks of EtOH OSA. To tag acquisition-recruited dMSNs, mice received 4-OHT after two consecutive acquisition sessions. Mice were then trained for 9 d of extinction and waited 2 weeks to allow virus expression. This was followed by a test extinction session with C21 (1 mg/kg, i.p.), 3 additional days of extinction, and finally a cue-induced reinstatement test with C21. ***E*,** During the last three FR3 acquisition sessions (gray shading indicates tagging sessions), active presses were similar among groups and significantly higher than inactive presses. Mixed-effect analysis: Group: *F*_(2,56)_ = 1.46, *p* = 0.24; Lever: *F*_(1,28)_ = 50.75, ^###^*p* < 0.001. ***F*,** Extinction training significantly reduced active presses to inactive level. Three-way ANOVA: Lever × Session: *F*_(8,224)_ = 15.86, ^###^*p* < 0.001. ***G*,** Chemogenetic inhibition of Acq-recruited dMSNs during extinction did not alter active-pressing (as normalized to that in the last extinction session). Mann-Whitney test: *U* = 88, *p* = 0.319. ***H*,** During cue-induced reinstatement, chemogenetic inhibition of Acq-recruited dMSNs suppressed active pressing. Only mCh group exhibited an increase of active pressing from the baseline. Compared to 100% theoretical mean: one-sample t test: mCh: *t*_(14)_ = 3.39, ^††^*p* = 0.004; Gi: *t*_(14)_ = 1.64, *p* = 0.124. Group comparison: Mann-Whitney test: *U* = 49, \*\**p* = 0.0075. ***I*,** After chemogenetic manipulation during cue-induced alcohol reinstatement, some mice were re-trained for sucrose (Suc) OSA in the same chamber. Mice underwent 5 d of FR3 sucrose training and 9 days of extinction before a sucrose-cue-induced reinstatement test with C21. ***J*,** Chemogenetic inhibition of alcohol acquisition-recruited dMSNs did not affect sucrose-cue-induced reinstatement. Both groups increased active pressing compared to their baselines. Compared to 100% theoretical mean: one-sample t test: mCh: *t*_(4)_ = 2.47, *p* = 0.069; Gi: *t*_(5)_ = 4.18, ^††^*p* = 0.009. Group comparison: unpaired t test: *t*_(9)_ = 1.01, *p* = 0.34. *K*, Summary diagram illustrating that acquisition-recruited dMSNs are specifically involved in driving alcohol relapse. n = 15 mice (E-H; mCh), 15 mice (E-H; Gi), 5 mice (J; mCh), 6 mice (J; Gi).

For selective chemogenetic manipulation of learning-recruited dMSNs, we employed a multi-layered strategy. First, we delivered a retrograde AAV (AAVretro-DIO-FLPo) into the SNr, leveraging the direct pathway. This method allowed dMSNs, but not other striatal neurons, to transport the AAV retrogradely to the DMS, resulting in selective DIO-FLPo presentation in dMSNs (Fig. 3B). Second, we utilized the ArcTRAP system to trigger CreER expression in neurons recruited during acquisition of alcohol learning. The introduction of 4-OHT facilitated translocation of CreER to the nuclei of activated neurons (Fig. 3B). Third, nuclear CreER elicited FLPo expression in acquisition-recruited dMSNs. FLPo then triggered FLPo-dependent hM4Di (Gi) or mCherry (mCh, as a control) expression exclusively in these acquisition-recruited dMSNs (Fig. 3B, 3C; Fig. S3A).

Following virus infusion, these ArcTRAP mice were trained for operant alcohol learning until they reached stable FR3 performance (Fig. 3D, 3E). Tagging was conducted during two consecutive FR3 sessions (Fig. 3D, 3E), followed by 9 d of extinction training in the same operant chamber (Fig. 3D, 3F). During an extinction memory test (no alcohol or cue), we found that chemogenetic inhibition of acquisition-recruited dMSNs (via C21 injection, 1 mg/kg)^50^ did not alter lever pressing (Fig. 3G; Fig. S3B, S3C), indicating that acquisition-recruited dMSNs might not be involved in extinction memory retrieval. After three additional days of extinction (Fig. 3D; Fig. S3D), mice underwent a cue-induced reinstatement test. The mCh group showed increased active presses relative to the final extinction day (Fig. 3H; S3E-S3G), reflecting successful alcohol memory retrieval. In contrast, chemogenetic inhibition of acquisition-tagged dMSNs significantly suppressed reinstatement of active pressing (Fig. 3H; Fig. S3E-S3G), demonstrating that these dMSNs are required for alcohol memory retrieval. Inactive lever presses were unaffected (Fig. S3B, S3E), and open field testing confirmed that this manipulation did not impair general locomotion (Fig. S3H). Together, these findings suggest that acquisition-recruited dMSNs specifically encode cue-alcohol memory without influencing general motor functions.

To determine whether these dMSNs selectively encode alcohol-associated memories, we next asked whether the same dMSNs are involved in the reinstatement of sucrose seeking. We retrained mice to self-administer sucrose pellets in the same operant chamber. The previous inactive lever for alcohol now became the active lever for sucrose, and vice versa (Fig. 3I). A distinct cue light was paired with sucrose delivery. Mice readily acquired this new paradigm (Fig. S3I). After 9 d of extinction, we used the sucrose-paired cue to test reinstatement. Critically, chemogenetic inhibition of alcohol acquisition-recruited dMSNs did not alter sucrose reinstatement, with both groups showing similar increases in active presses (Fig. 3J; Fig. S3J). This indicates that these alcohol acquisition-recruited neurons do not generalize to other reward-associated memories.

Lastly, we examined whether dMSNs activated in the home-cage, reflecting baseline activity, could drive alcohol relapse. A new group of mice were trained for operant alcohol learning (Fig. S4A, S4B), and dMSNs were tagged in their home-cages (∼24 h after the last acquisition session) with Gi. Following 9 d of extinction training (Fig. S4C), a cue-induced reinstatement test was conducted. Notably, inhibiting home-cage-activated dMSNs did not alter active pressing during reinstatement (Fig. S4D), suggesting that baseline-activated dMSNs did not control alcohol relapse.

In summary, these findings demonstrate that alcohol acquisition-recruited dMSNs are both necessary and selectively engaged during the retrieval of cue-alcohol memories underlying relapse (Fig. 3K). Thus, acquisition-recruited dMSNs constitute a distinct striatal engram population that specifically encodes alcohol-associated memories and drives relapse behavior.

### Alcohol extinction-recruited dMSNs selectively drive alcohol, but not sucrose, extinction memory and suppress alcohol relapse

Having established that acquisition-recruited dMSNs are required for retrieving alcohol-associated memories that drive relapse, we next asked whether extinction-recruited dMSNs function as an opposing engram. Specifically, we hypothesized that activating extinction-recruited dMSNs would promote extinction memory retrieval and actively suppress alcohol relapse (Fig. 4A).

**Figure 4.**
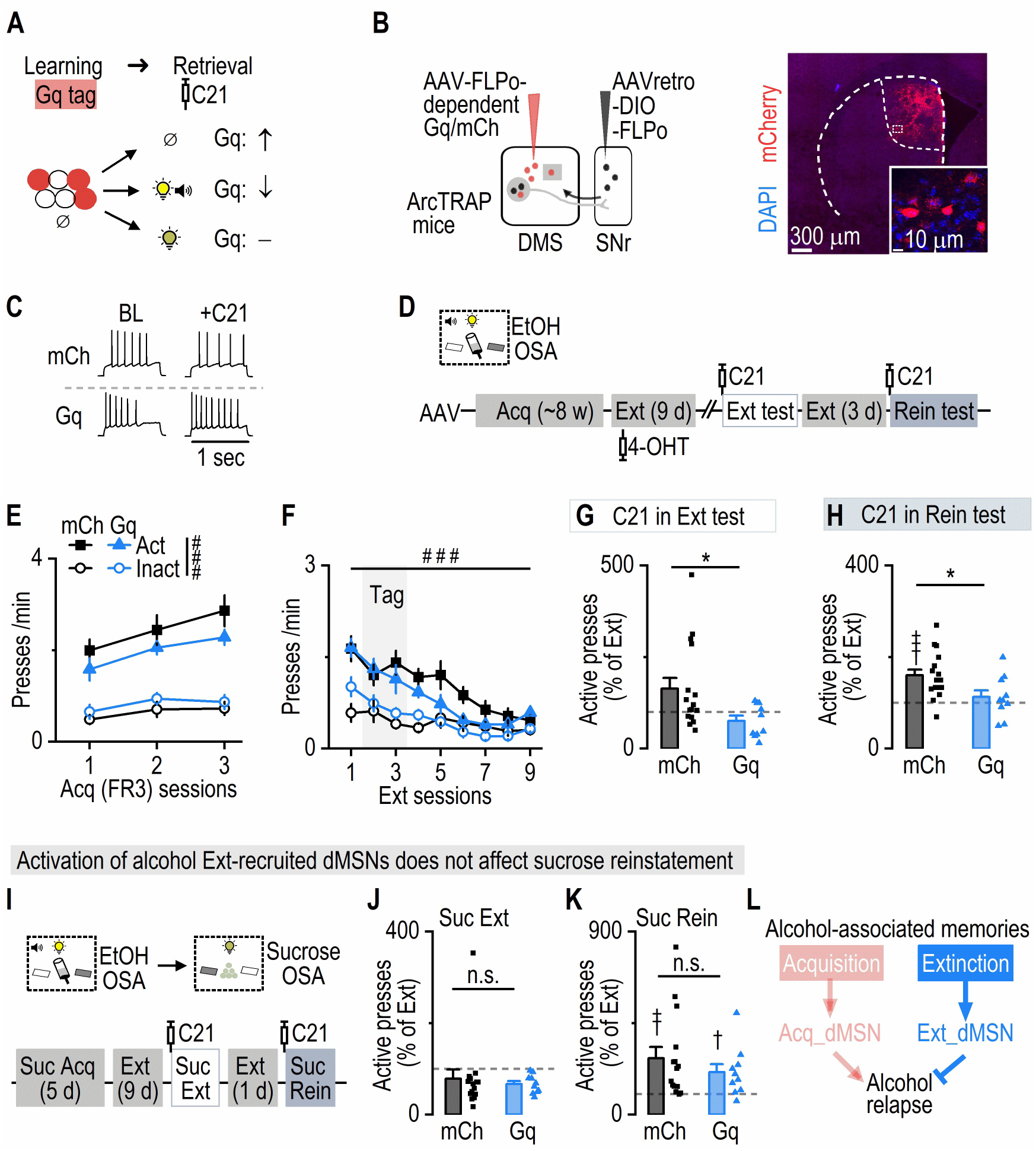
Chemogenetic activation of extinction-recruited dMSNs drives extinction and suppresses the reinstatement of alcohol, but not sucrose, seeking. ***A*,** Schematic illustrating the hypothesis: chemogenetic excitation of extinction-tagged neurons (expressing Gq) promotes extinction memory retrieval (top) and suppresses the retrieval of the cue-alcohol memory (middle), while leaving other memories unaffected (bottom). ***B*,** Schematic of dual-virus infusions in ArcTRAP mice to direct hM3Dq or mCherry expression to extinction-recruited dMSNs. AAVretro-DIO-FLPo was infused into the SNr, followed by FLPo-dependent DREADDs (AAV-fDIO-hM3Dq-mCherry [Gq] or AAV-fDIO-mCherry [mCh]) in the DMS. In the DMS, circles represent dMSNs and squares represent non-dMSNs. ***C*,** Electrophysiological recordings demonstrate that C21 did not alter evoked firing rates in mCh-tagged neurons but increased evoked firing rates in Gq-tagged neurons under identical current injections. ***D*,** Experimental timeline. Mice were trained in EtOH OSA (∼8 weeks), then received 9 d of extinction. To tag extinction-recruited dMSNs, 4-OHT was administered during the 2nd and 3rd extinction sessions. Two weeks later, mice underwent an extinction test with C21 (1 mg/kg, i.p.), followed by 3 more extinction days and a final cue-induced reinstatement test with C21. ***E*,** Both mCh and Gq groups showed similar active-press levels in the last 3 d of OSA, which exceeded their inactive-press levels. Three-way ANOVA: Group: *F*_(1,24)_ = 0.42, *p* = 0.523; Lever: *F*_(1,24)_ = 65.28, ^###^*p* < 0.001. ***F*,** During extinction, active presses decreased to the inactive-level in mCh and Gq groups. Lever × Session: *F*_(8,192)_ = 8.56, ^###^*p* < 0.001. ***G*,** During extinction test, activating extinction-recruited dMSNs reduced normalized active presses relative to the last extinction session compared to mCh. Mann-Whitney test: *U* = 39.50, \**p* = 0.032. ***H*,** Chemogenetic activation of extinction-recruited dMSNs suppressed active presses during reinstatement. Compared to 100% theoretical mean: one-sample t test: mCh: *t*_(15)_ = 4.69, ^†††^*p* < 0.001; Gq: *t*_(9)_ = 0.85, *p* = 0.42. Group comparison: unpaired t test: *t*_(24)_ = 2.39, \**p* = 0.025. ***I*,** After chemogenetic manipulation during cue-induced alcohol reinstatement, mice were re-trained for sucrose OSA in the same chamber. Mice underwent 5 d of FR3 sucrose training and 9 days of extinction before an extinction test with C21. This was followed by another extinction and a sucrose-cue-induced reinstatement test with C21. ***J*,** Chemogenetic activation of alcohol extinction-recruited dMSNs did not affect the extinction of sucrose memory. Mann-Whitny test: *U* = 65, *p* = 0.59. *K,* Chemogenetic activation of alcohol extinction-recruited dMSNs did not alter sucrose-cue-induced reinstatement; both groups exhibited similar increase in active presses. Compared to 100% theoretical mean: one-sample t test: mCh: *t*_(14)_ = 3.28, ^††^*p* = 0.006; Gq: *t*_(9)_ = 2.81, ^†^*p* = 0.021. Group comparison: Mann-Whitny test: *U* = 64, *p* = 0.57. ***L*,** Summary diagram illustrates that extinction-recruited dMSNs specifically support retrieval of alcohol extinction memory and their activation suppresses alcohol relapse. n = 16 mice (mCh; E-H), 10 mice (Gq; E-H), 15 mice (mCh; J, K), and 10 mice (Gq; J, K).

To test this, we selectively tagged extinction-recruited dMSNs with hM3Dq (or mCh as a control) in ArcTRAP mice (Fig. 4B, 4C). Animals underwent operant alcohol self-administration followed by 9 days of extinction, with neuronal tagging performed during the second and third extinction sessions (Fig. 4D). Both groups displayed comparable acquisition and extinction performance (Fig. 4E, 4F). Three weeks after extinction training, we tested the causal role of extinction-recruited dMSNs in an extinction memory retrieval test. Notably, chemogenetic activation of extinction-recruited dMSNs significantly reduced active lever presses compared to the mCh control group (Fig. 4G; Fig. S5A). These results indicate that extinction-recruited dMSNs were sufficient to facilitate extinction memory retrieval. After additional 3-d extinction training (Fig. S5B), we examined the effects of manipulating extinction-recruited dMSNs during cue-induced reinstatement. Importantly, chemogenetic activation of extinction-recruited dMSNs significantly reduced active lever pressing during reinstatement, without affecting inactive lever pressing (Fig. 4H; Fig. S5C) or general locomotor activity measured in an open-field test (Fig. S5D). Thus, activation of extinction-recruited dMSNs actively suppressed relapse, indicating their critical role as a memory-specific inhibitory engram population.

Lastly, we tested whether extinction-recruited dMSNs selectively encode alcohol-specific extinction memories, or whether they might generalize to extinction of other reward-associated memories. To do so, we retrained the same mice to self-administer sucrose pellets in the same operant chambers, with the previously inactive lever now serving as the active lever for sucrose, accompanied by a distinct cue (Fig. 4I). Both groups of mice readily learned the sucrose task and successfully underwent 9-d of extinction training (Fig. S5E). Crucially, chemogenetic activation of the alcohol extinction-recruited dMSNs did not alter active or inactive lever presses during sucrose extinction (Fig. 4J; Fig. S5F). Moreover, activation of these neurons did not influence reinstatement behavior triggered by sucrose-associated cues, with both groups showing similar increases in active lever pressing and no differences in inactive lever presses (Fig. 4K; Fig. S5G).

Together, these findings demonstrate that extinction-recruited dMSNs constitute a specialized striatal engram selectively encoding alcohol extinction memories. Reactivation of this engram actively facilitates extinction memory retrieval and suppresses relapse to alcohol seeking without generalizing to natural rewards such as sucrose (Fig. 4L). These findings, together with our prior results, support a dual-engram architecture in the dorsal striatum in which alcohol acquisition and extinction recruit anatomically and functionally distinct populations of dMSNs. Each population encodes a behaviorally specific memory, either promoting or suppressing alcohol relapse.

### Alcohol acquisition induces persistent potentiation of mPFC–dMSN engram synapses

Memory formation is typically linked to synaptic plasticity^10,13,51^. Specifically, it has been proposed that synaptic strength at engram-specific synapses constitutes the cellular substrate of memory^52–55^. Having established that distinct engram populations of DMS dMSNs encode acquisition versus extinction of alcohol-associated memories, we next sought to identify where and how alcohol-related memories are physically encoded within these learning-recruited dMSNs. A defining feature of engram cells is that they undergo learning-induced synaptic modifications at specific input pathways, thereby storing long-lasting memory traces^12,13^. Given that acquisition-recruited dMSNs specifically encoded cue-alcohol memories and directly controlled relapse behavior (Figs. 1-3), we reasoned that acquisition-related synaptic plasticity onto these dMSNs might represent a critical substrate underlying alcohol memory encoding. In particular, the medial prefrontal cortex (mPFC) provides crucial excitatory input to dMSNs implicated in alcohol-seeking behaviors^56,57^; thus, we hypothesized that synaptic transmission between alcohol-activated mPFC neurons and acquisition-recruited dMSNs may exhibit selective potentiation following alcohol acquisition, representing the physical engram trace (Fig. 5A). To test this hypothesis, we performed input-specific electrophysiological recordings comparing synaptic strength of alcohol-activated mPFC neurons (mPFC^tag^) onto acquisition-recruited versus non-recruited dMSNs (dMSN^tag^ versus dMSN^non-tag^, respectively).

**Figure 5.**
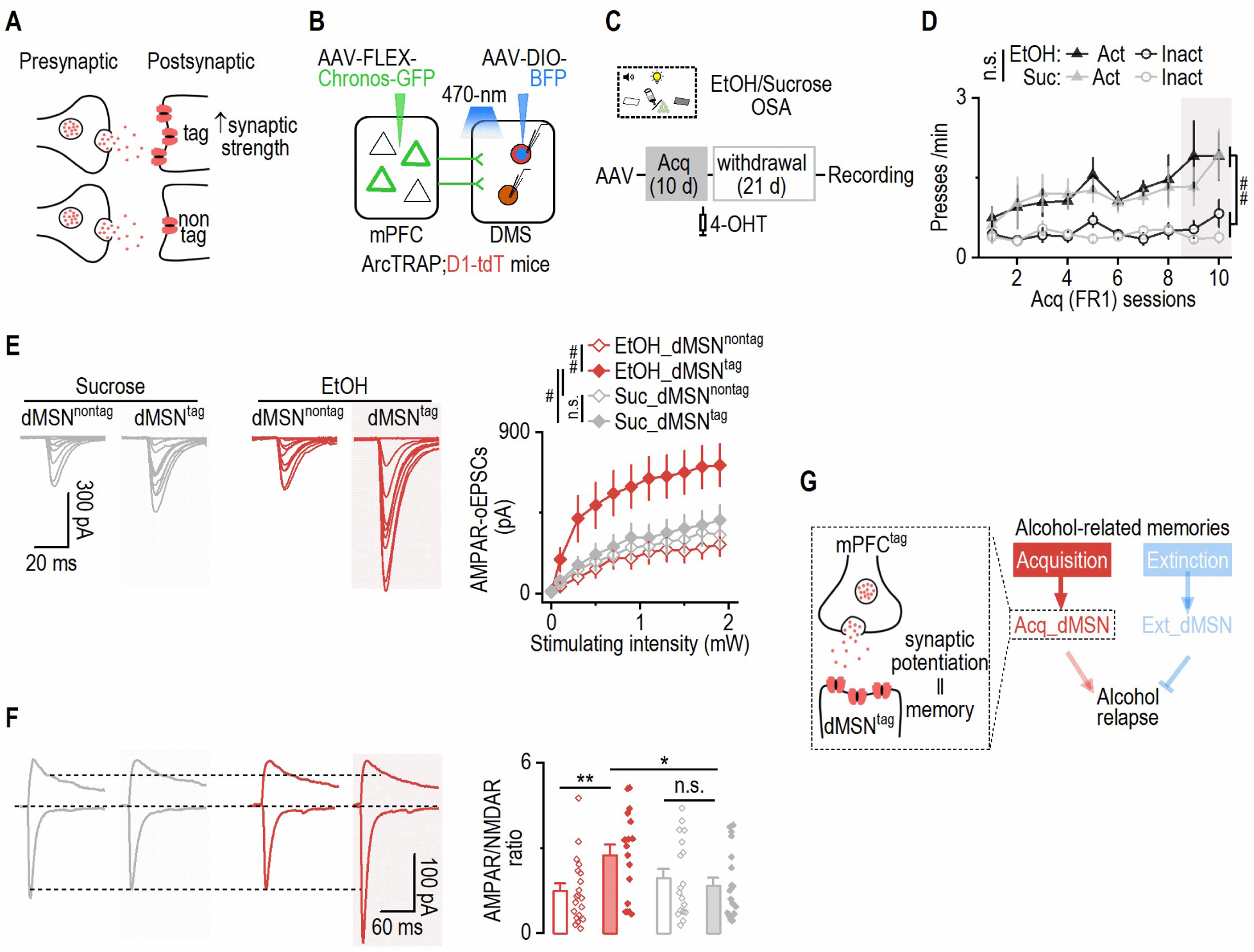
Alcohol acquisition-recruited mPFC-to-dMSN synapses exhibit preferential and long-lasting potentiation. ***A*,** Schematic illustrating the hypothesis: operant learning induces preferential synaptic potentiation between presynaptically tagged neurons and postsynaptically tagged neurons, measurable via electrophysiology. ***B*,** Schematic of virus infusion and recording. AAV-FLEX-Chronos-GFP was infused into the mPFC, and AAV-DIO-BFP (blue fluorescent protein) was infused into the DMS of ArcTRAP;D1-tdT mice. Optically induced excitatory postsynaptic currents (oEPSCs) were measured by stimulating Chronos-expressing fibers from mPFC^tag^ to dMSN^tag^ or _dMSNnon-tag._ ***C*,** Following virus infusion, mice underwent 10 d of EtOH or Sucrose (Suc) OSA training under an FR1 schedule. 4-OHT was administered after the last two sessions to tag activated neurons. Mice then underwent 21 d withdrawal in home-cages before electrophysiology. ***D*,** Mice trained for alcohol and sucrose had similar active presses throughout training, both significantly higher than inactive presses. Three-way ANOVA: Group: *F*_(1,8)_ = 0.14, *p* = 0.72; Lever: *F*_(1,18)_ = 16.13, ^##^*p* = 0.004. ***E*,** After 21-d withdrawal, AMPAR-oEPSCs from mPFC^tag^ inputs were higher in dMSN^tag^ than dMSN^non-tag^ in the alcohol group; this difference was not seen in the sucrose group. Mixed-effects analysis: Group: *F*_(3, 78)_ = 4.86, *p* = 0.0038. Tukey’s multiple comparisons test: EtOH-dMSN^non-tag^ vs. EtOH_dMSN^tag^: *q* = 4.93, ^##^*p* = 0.0044; EtOH-dMSN^tag^ vs. Suc_dMSN^tag^: *q* = 4.27, ^#^*p* = 0.018; EtOH-dMSN^tag^ vs. Suc_dMSN^non-tag^: *q* = 3.73, ^#^*p* = 0.048; Suc-dMSN^non-tag^ vs. Suc_dMSN^tag^: *q* = 0.59, *p* = 0.97. ***F*,** AMPAR/NMDAR ratios were strongest in dMSN^tag^ compared to dMSN^non-tag^ in the alcohol group, and also higher than dMSN^tag^ in the sucrose group. Two-way ANOVA: Group × Tag: *F*_(1, 70)_ = 6.34, *p* = 0.014. Sidak’s multiple comparisons test: dMSN^non-tag^ vs. dMSN^tag^ in EtOH: *t* = 2.90, \*\**p* = 0.01; dMSN^non-tag^ vs. dMSN^tag^ in Suc: *t* = 0.66, *p* = 0.76; EtOH vs. Suc for dMSN^tag^: *t* = 2.47, \**p* = 0.031. ***G*,** Summary schematic illustrating that relapsing-driving alcohol-associated memories encoded by acquisition-recruited dMSNs are likely physically stored via synaptic potentiation at mPFC^tag^-to-dMSN^tag^ synapses. n = 5 mice per group (D), 21 neurons from 5 mice (21/5; EtOH-dMSN^non-tag^; E), 20/5 (EtOH-dMSN^tag^; E), 20/5 (Suc-dMSN^non-tag^; E), 21/5 (Suc-dMSN^tag^; E), 20/5 (EtOH-dMSN^non-tag^; F), 17/5 (EtOH-dMSN^tag^; F), 18/5 (Suc-dMSN^non-tag^; F), and 19/5 (Suc-dMSN^tag^; F).

First, we selectively expressed Chrimson-tdT in mPFC^tag^ of triple transgenic mice (ArcTRAP;Ai32 [ChR2-EYFP];D1-tdT) (Fig. S6A). Tagging was conducted during two consecutive acquisition sessions, and slice electrophysiology was performed three weeks later (Fig. S6B, S6C). dMSN^tag^ were identified by tdT and EYFP expression and verified by sustained ChR2-mediated photocurrents during light stimulation (Fig. S6D, S6E). Using 590-nm light to stimulate mPFC^tag^ axons in the DMS (Fig. S6F), we found that mPFC^tag^ inputs onto dMSN^tag^ exhibited significantly larger AMPA receptor-mediated optically evoked excitatory postsynaptic currents (EPSCs) and higher AMPAR/NMDAR ratios, as compared to inputs onto dMSN^non-tag^ (Fig. S6G, S6H). These changes in synaptic strength were likely postsynaptic, as no differences in paired-pulse ratios were observed (Fig. S6I)^58,59^. These results suggest that acquisition of operant alcohol learning promotes the formation of stronger synaptic connections between behaviorally defined cortical inputs and engram dMSNs.

Neurons recruited by natural rewards have also been shown to exhibit synaptic strengthening, as compared to surrounding non-recruited neurons^60^. However, prior work suggests that synaptic plasticity induced by natural rewards may be less enduring than that induced by addictive substances^61,62^. We therefore next asked whether mPFC^tag^-dMSN^tag^ synaptic potentiation observed after alcohol acquisition persisted longer than that induced by sucrose learning. To investigate this, we expressed Chronos-GFP in mPFC^tag^ of ArcTRAP;D1-tdT mice and trained them for either alcohol or sucrose operant learning (10 d), with neuronal tagging performed during the final two acquisition sessions (Fig. 5B-5D). Following training, animals were withdrawn to their home cages for 21 d to assess the persistence of synaptic changes (Fig. 5C). Although spontaneous EPSCs were similar between groups (Fig. S7A), both AMPAR-mediated optically induced EPSC amplitudes and AMPAR/NMDAR ratios remained significantly higher in dMSN^tag^ than in dMSN^non-tag^ in the alcohol group, but not in the sucrose group (Fig. 5E, 5F). As with prior experiments, these effects were not associated with differences in paired-pulse ratio, indicating a postsynaptic locus of plasticity (Fig. S7B).

Taken together, these data demonstrate that alcohol acquisition induces robust and long-lasting potentiation of corticostriatal synapses between alcohol-activated mPFC neurons and recruited dMSNs. These input-specific and persistent changes are consistent with a synaptic engram model, in which memory traces are stored at strengthened synapses linking behaviorally defined presynaptic and postsynaptic engram populations (Fig. 5G).

### Optogenetic strengthening of mPFC-to-dMSN synapses reconstitutes a functional engram sufficient to drive relapse-like behavior

The preceding experiments revealed that alcohol acquisition strengthens synaptic transmission between alcohol-activated mPFC neurons and acquisition-recruited dMSNs, supporting the idea that memory traces are stored at behaviorally defined corticostriatal synapses. To determine whether this synaptic potentiation alone was sufficient to recreate a functional memory engram capable of driving relapse-like behavior, we employed oICSS^25,34,63,64^ to mimic alcohol-induced strengthening at mPFC-to-dMSN synapses (Fig. 6A). This approach enabled us to isolate the role of synaptic potentiation from other alcohol-related changes and assess whether artificial strengthening of this input-specific corticostriatal pathway could recreate a functional memory engram sufficient to support relapse-like behavior, independent of prior alcohol experience.

**Figure 6.**
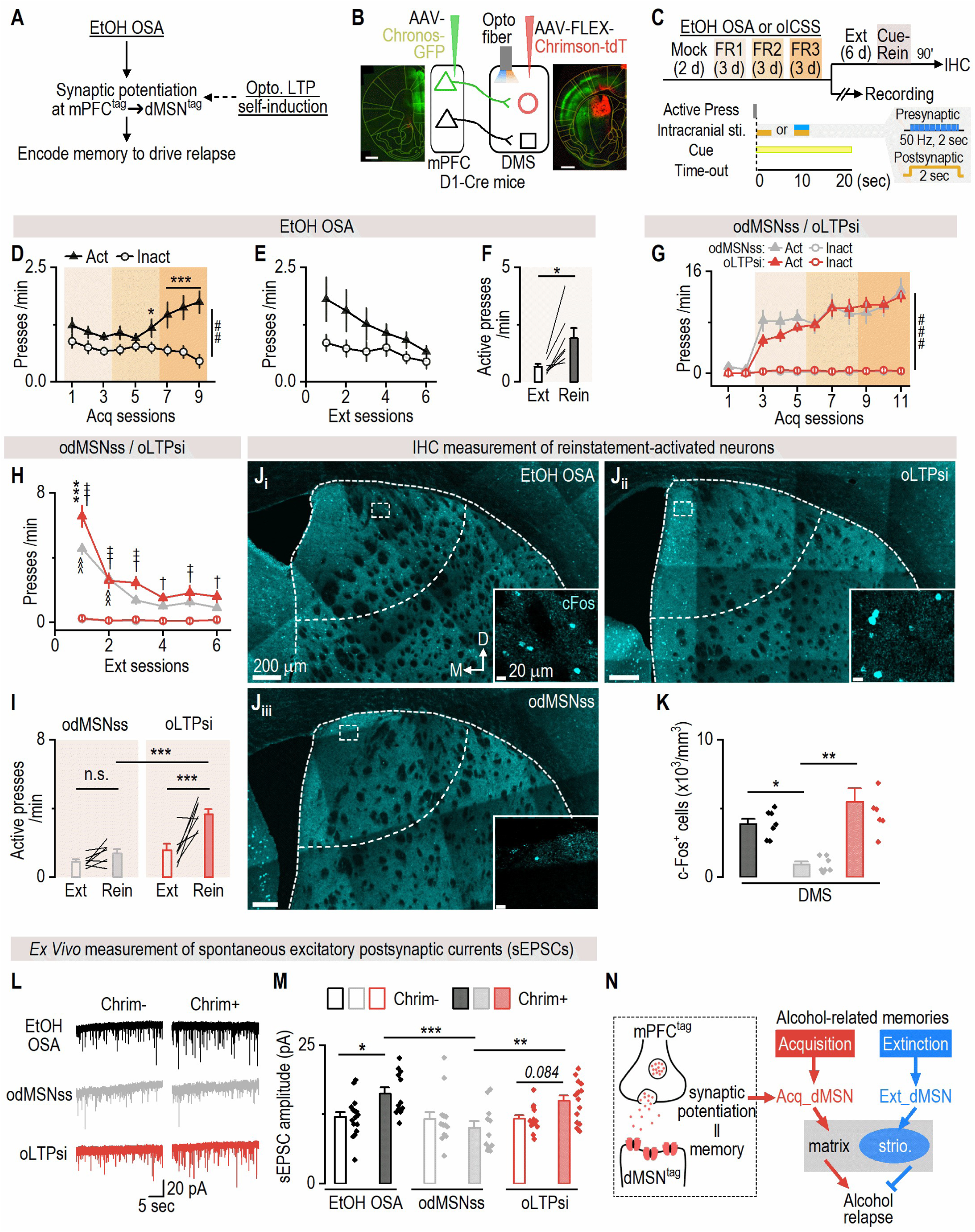
Optical self-induction of mPFC-to-dMSN synaptic potentiation is sufficient to form an alcohol-like cue memory to drive relapse-like behavior. ***A*,** Schematic illustrating the hypothesis: If EtOH OSA-induced synaptic potentiation between activated mPFC-dMSNs is the engram that encodes alcohol-related memories to drive relapse, optical self-induction of mPFC-to-dMSN synaptic potentiation (i.e., long-term potentiation [LTP]) should mimic the effect of EtOH OSA to encode a rewarding operant memory that can be retrieved by cue. ***B*,** The schematic of virus infusion and fiber implantation in D1-Cre mice. Sample images showing Chronos-GFP expression in the mPFC and Chrimson-tdT expression in the DMS. Scale bars: 600 µm (left), 800 µm (right). In the DMS, circle represents dMSN and square represents non-dMSN. ***C*,** Mice were separated into three groups. One group went through actual EtOH OSA for 9 d, while the other two groups were trained for oICSS using either optical LTP self-induction (oLTPsi) or optical dMSN self-stimulation (odMSNss, as a control) for 9 d. After the last day of training, some mice underwent 6 d of extinction, followed by a cue-induced reinstatement test and were then sacrificed for c-Fos IHC. The rest of the mice were sacrificed for electrophysiological recordings 1-3 d after the last training. Bottom right: The schematic of the oICSS protocol. ***D*,** In the EtOH OSA group, active presses gradually increased over inactive presses across the training period. Two-way RM ANOVA: Lever: *F*_(1,10)_ = 25.27, ^###^*p* < 0.001. Sidak’s multiple comparisons test: Act vs. Inact: sessions 1-6, *t* = 2.42, 2.31, 2.16, 2.59, 1.21, 3.01, 4.31, 6.79, 9.04; \**p* = 0.031 for session 6 and \*\*\**p* < 0.001 for sessions 7-9. ***E*,** In the EtOH OSA group, extinction training gradually decreased the active presses to the inactive level. Two-way RM ANOVA: Lever × Session: *F*_(5,30)_ = 1.52, *p* = 0.21. Sidak’s multiple comparisons test: Act vs. Inact: sessions 1-6, *t* = 3.97, 3.45, 2.51, 1.32, 1.59, 0.90; \*\**p* = 0.0025 for session 1, \**p* = 0.01 for session 2, and *p* > 0.05 for sessions 3-6. ***F*,** During alcohol cue-induced reinstatement, active presses were significantly increased compared to the last extinction. Paired t test: *t*_(6)_ = 3.58, \**p* = 0.012. ***G*,** In both oICSS groups, active presses significantly exceeded inactive presses throughout training, with no difference between groups. Three-way ANOVA: Lever: *F*_(1,18)_ = 106.7, ^###^*p* < 0.001; oLTPsi vs. odMSNss: *F*_(1,18)_ = 0.15, *p* = 0.71 ***H*,** During extinction training, the oLTPsi group had more active presses overall than the odMSNss group (Three-way ANOVA: Group: *F*_(1, 12)_ = 5.12, *p* = 0.043; Act in oLTPsi vs. odMSNss in session 1: *q* = 7.57, \*\*\**p* < 0.001). In the oLTPsi group, active presses remained consistently higher than inactive presses (Act vs. Inact in oLTPsi from sessions 1-6: *q* = 24.41, 9.56, 8.86, 5.55, 6.77, 5.44, ^†††^*p* < 0.001 for session 1-3, ^†^*p* < 0.05 for sessions 4 and 6, ^††^*p* < 0.01 for session 5), whereas active presses in the odMSNss group quickly decreased to the inactive level (Act vs. Inact in odMSNss from sessions 1-6: *q* = 16.90, 10.00, 4.94, 3.56, 4.34, 3.07, ^^^^^*p* < 0.001 for sessions 1-2 and *p* > 0.05 for sessions 3-6). ***I*,** During oICSS cue-induced reinstatement, the oLTPsi group showed a significant increase in active presses compared to their last extinction session, while the odMSNss group did not. Two-way RM ANOVA: Group × Session: *F*_(1,12)_ = 7.70, *p* = 0.017. Sidak’s multiple comparisons test: Ext vs. Rein: *t* = 1.22, *p* = 0.43 (odMSNss); *t* = 5.14, \*\*\**p* < 0.001 (oLTPsi); odMSNss vs. oLTPsi in Rein: *t* = 5.92, \*\*\**p* < 0.001. *J_i-iii_*, Sample images of IHC showing increased c-Fos expression in the EtOH OSA (i) and oLTPsi (ii) groups compared to the odMSNss (iii) group in the striatal area after cue-induced reinstatement. ***K*,** More c-Fos^+^ cells were observed in the DMS of the EtOH OSA and oLTPsi groups relative to the odMSNss group. Kruskal-Wallis test: *H*_(2)_ *=* 14.04, *p* < 0.001. Dunn’s multiple comparisons test: EtOH vs. odMSNss: *z* = 2.76, \**p* = 0.018; oLTPsi vs. odMSNss: *z* = 3.58, \*\**p* = 0.001. ***L*,** Sample traces of spontaneous excitatory postsynaptic currents (sEPSCs) recorded from Chrimson^-^ (Chrim^-^) and Chrimson^+^ (Chrim^+^) neurons in all three groups. ***M*,** sEPSC amplitudes were greater in Chrim^+^ than in Chrim^−^ neurons in the EtOH OSA and oLTPsi groups, but not in the odMSNss group. Moreover, sEPSC amplitudes in Chrim^+^ neurons of EtOH OSA and oLTPsi groups were higher than those in the odMSNss group. Two-way ANOVA: Group × Neuron (Chrim^-^ vs. Chrim^+^): *F*_(2,73)_ = 5.21, *p* = 0.0077. Sidak’s multiple comparisons test: Chrim^-^ vs Chrim^+^: *t* = 2.90, \**p* = 0.015 (EtOH OSA); *t* = 1.04, *p* = 0.66 (odMSNss); *t* = 2.23, *p* = 0.084 (oLTPsi). Chrim^+^ in EtOH OSA vs. odMSNss: *t* = 3.94, \*\*\**p* < 0.001; Chrim^+^ in odMSNss vs. oLTPsi: *t* = 3.22, \*\**p* = 0.0057. *N*, Schematic summary of key findings. Artificial corticostriatal potentiation functionally mimics the naturally formed acquisition engrams, which are preferentially localized in the striatal matrix and drives alcohol relapse. In contrast, extinction engrams are preferentially localized in the striosome and suppress relapse. n = 11 mice (D), 7 mice (E, F), 10 mice per group (G), 7 mice per group (H-I, K), 16 neurons/4 mice (16/4; Chrim^-^ in EtOH; M), 12/4 (Chrim^+^ in EtOH; M), 13/3 (Chrim^-^ in odMSNss; M), 11/3 (Chrim^+^ in odMSNss; M), 13/3 (Chrim^-^ in oLTPsi; M), 14/3 (Chrim^+^ in oLTPsi; M).

Previous studies have shown that optogenetic dMSN self-stimulation (odMSNss) is instrumentally reinforcing but does not form a conditioned association with paired cues^25,63,65^. In contrast, the mPFC is essential for encoding cue-related information^66,67^. We therefore postulated that animals trained to lever-press to self-induce optical mPFC-to-dMSN long-term potentiation (oLTPsi; paired with discrete cues) would form a lasting cue memory that could reinstate seeking behavior, while odMSNss alone would not. To test this, we utilized a dual-channel optogenetic approach^56,57,68,69^, expressing Chronos (activated by 473-nm light) in the mPFC and Chrimson (activated by 590-nm light) in the DMS dMSNs of D1-Cre mice (Fig. 6B, 6C). We additionally verified that each opsin was selectively activated by its designated wavelength, as stimulating single-opsin mice with the non-matching wavelength failed to produce operant responding (Fig. S8A-S8B). To train mice for odMSNss, we tested various parameters and found that a 2-sec constant 590-nm stimulation delivered immediately after an active press induced the strongest operant reinforcement (Fig. S8C-S8F). Thus, this protocol was subsequently used for training. To train mice for oLTPsi, a presynaptic mPFC stimulation protocol was paired with the odMSNss protocol and delivered simultaneously^56,57,68,69^ after the active press (Fig. 6C) as such pairing is known to induce corticostriatal potentiation^57^. Importantly, active presses during oICSS triggered optical stimulation and cue presentation (light+tone). As a reference, we also included a group that underwent conventional alcohol operant learning to allow for behavioral and physiological comparisons between artificial oICSS and real alcohol learning.

Mice trained for alcohol operant learning showed a progressive increase in active presses over inactive ones; this behavior was subsequently extinguished, but reinstated by alcohol-paired cues (Fig. 6D-6F). Both odMSNss and oLTPsi groups also exhibited robust acquisition of active presses (Fig. 6G)^25,63^. However, the oLTPsi group exhibited a delayed extinction compared to the odMSNss group (Fig. 6H), suggesting that a more persistent operant memory had been encoded by oLTPsi. Crucially, stimulation-paired cues only reinstated active presses in the oLTPsi group, but not in the odMSNss group (Fig. 6I), indicating successful encoding of a cue memory during mPFC-to-dMSN synaptic potentiation, mirroring alcohol relapse (Fig. 6F). During the reinstatement test, c-Fos staining revealed higher dorsal striatal activity (c-Fos^+^ cell density) in both the alcohol and oLTPsi groups compared to the odMSNss group (Fig. 6J-6K). Additionally, we measured glutamatergic synaptic transmission in activated dMSNs (Chrimson^+^ neurons) and surrounding nonactivated neurons (Chrimson^-^ neurons). The amplitude of spontaneous EPSCs—but not their frequency—was significantly higher in activated dMSNs in the alcohol and oLTPsi groups, but not in the odMSNss group (Fig. 6L, 6M; Fig. S8G). Although non-activated dMSNs in the oLTPsi group also received mPFC inputs, they were not expected to undergo LTP as prior work has shown that mPFC stimulation alone does not induce corticostriatal potentiation^56,57^. Importantly, similar findings were replicated in D1-Cre rats (Fig. S9), confirming cross-species validity.

In summary, these findings demonstrate that artificial potentiation at mPFC-to-dMSN synapses is sufficient to recreate a functional acquisition engram that mimics alcohol memory and drives relapse-like behavior. This optogenetically-induced memory recapitulates key features of alcohol learning, including cue-induced reinstatement, increased striatal activation, and synaptic strengthening in activated dMSNs. These results establish that input-specific corticostriatal plasticity is not only a correlate but a causal substrate of relapse memory, underscoring the central role of mPFC-to-dMSN engram synapses in encoding alcohol-associated memories (Fig. 6N).

## Discussion

Our findings overturn the prevailing view that opposing behavioral memories arise from different striatal pathways (i.e., dMSN versus indirect-pathway MSN). We show that both relapse-promoting and relapse-suppressing alcohol-associated memories are encoded within the same genetically defined striatal cell type, dMSNs, but in anatomically and functionally distinct ensembles. Acquisition of alcohol seeking engages matrix-enriched dMSNs that store cue-alcohol memories and drive relapse, whereas extinction recruits a striosome-enriched ensemble that stores extinction memories and suppresses relapse. We further identify the physical substrate of the relapse engram within input-specific synaptic potentiation between alcohol-activated mPFC inputs and acquisition-recruited dMSNs. Optogenetically mimicking this plasticity is sufficient to encode a persistent cue memory that triggers relapse-like behavior.

These results parallel recent hippocampal work showing that fear memory and its extinction are supported by distinct engrams^42^, but reveal a surprising twist. In the dorsal striatum, these opposing memories are encoded within the same genetically defined cell type, segregated by compartmental localization and circuit connectivity. This establishes the dual-engram principle in instrumental drug seeking and challenges the assumption that opposing memories require different neuronal types. Rather than overwriting the original memory, extinction emerges as an active learning process with its own dedicated engram, stored in parallel with the acquisition memory and competing for retrieval to control behavior. The biased localization of these engrams in matrix versus striosome compartments provides a structural basis for their divergent roles, consistent with the opposing behavioral tendencies of these compartments.

Why can dMSNs being recruited into alcohol-related engram ensembles encode instrumental memories? In the striatum, MSN recruitment is shaped by glutamatergic inputs and dopamine signaling^26^. Addictive substances elevate striatal dopamine^45,70^, enhancing glutamate-driven firing in dMSNs via both pre- and postsynaptic mechanisms, which may bias them toward engram allocation^71,72^. Consistent with this, several studies have shown that addictive substances preferentially activate dMSNs, but not other striatal cells^27,29^.

How extinction-recruited dMSNs suppress the expression of acquisition memories remains an open question. One possibility is via dopaminergic modulation. Extinction-recruited striosomal dMSNs strongly inhibit substantia nigra dopamine neurons, potentially reducing dopamine release during residual attempts to seek alcohol during early phases of extinction learning and thereby promoting LTD at acquisition engram synapses^73^. Such a mechanism could gradually weaken the original memory trace and bias retrieval toward extinction^55^.

These findings challenge the canonical view of the striatal direct pathway as a uniform “Go” circuit for motivated behavior^45^. Even within dMSNs, behavioral function is shaped not only by genetic identity but also by anatomical context and learning history, producing experience-defined ensembles with opposing roles. This framework highlights the need to study behaviorally recruited ensembles within genetically defined populations, rather than assuming functional uniformity based on cell type alone.

Finally, our results have direct translational implications. Extinction alone reduces relapse relative to abstinence, but targeted inhibition of the relapse engram or activation of the extinction engram further suppresses relapse. This suggest that combining behavioral therapy with ensemble-specific interventions could enhance treatment efficacy. By identifying the cellular and synaptic substrates of opposing alcohol-associated memories, our study provides a blueprint for precision targeting of maladaptive memories, with potential relevance to a broad range of neuropsychiatric disorders driven by persistent and competing memories.

## Supplementary Figures and Figure Legends

**Supplementary Figure 1.**
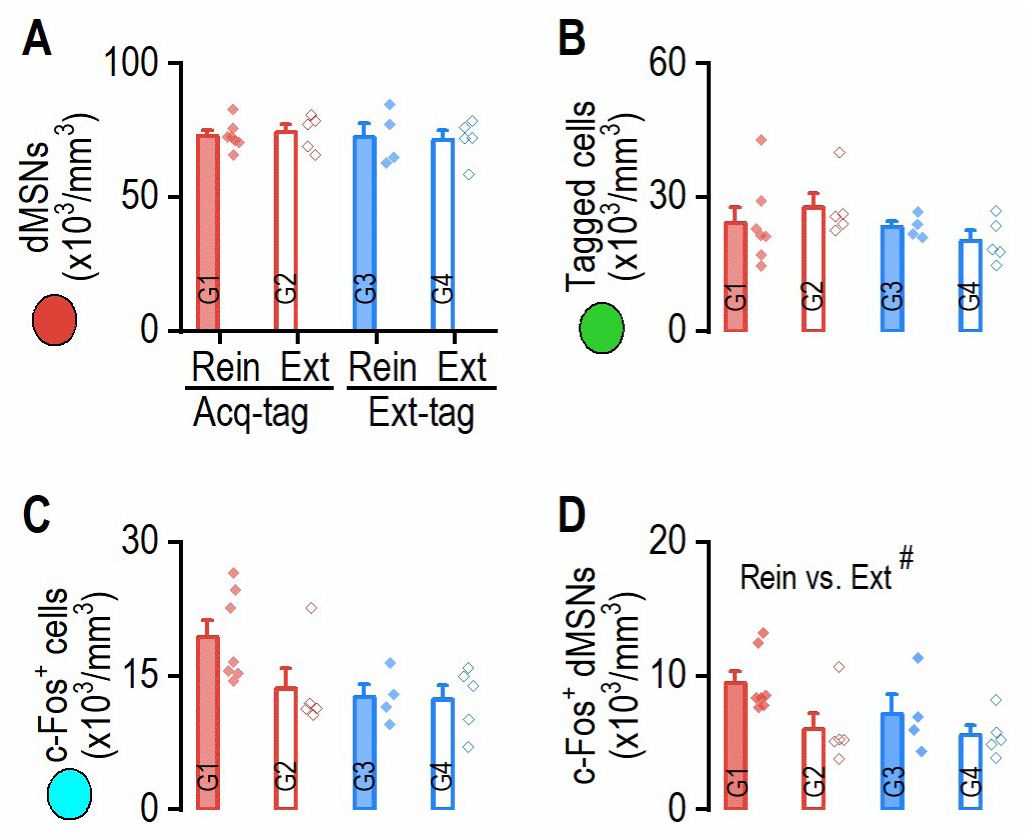
Histological analysis of tagged dMSNs and c-Fos^+^ dMSNs in acquisition and extinction-tagged groups after extinction or reinstatement test (Related to Figure 1) ***A*,** Density of dMSNs showed no difference among groups. Two-way ANOVA: Group × Test: *F*_(1, 17)_ = 0.13, *p* = 0.73. ***B*,** Density of tagged cells had no difference among groups. Two-way ANOVA: Group × Test: *F*_(1, 17)_ = 1.12, *p* = 0.31. ***C*,** Density of c-Fos^+^ cells had no difference among groups. Two-way ANOVA: Group × Test: *F*_(1, 17)_ = 1.95, *p* = 0.18. ***D*,** Groups tested for cue-induced reinstatement showed a higher density of c-Fos^+^ dMSNs compared to groups tested for extinction. Two-way ANOVA: Test (Rein vs Ext): *F*_(1, 17)_ = 5.52, ^#^*p* = 0.031. n = 7 mice (Acq-tag, Rein test), 5 mice (Acq-tag, Ext test), 4 mice (Ext-tag, Rein test), and 5 mice (Ext-tag, Ext test).

**Supplementary Figure 2.**
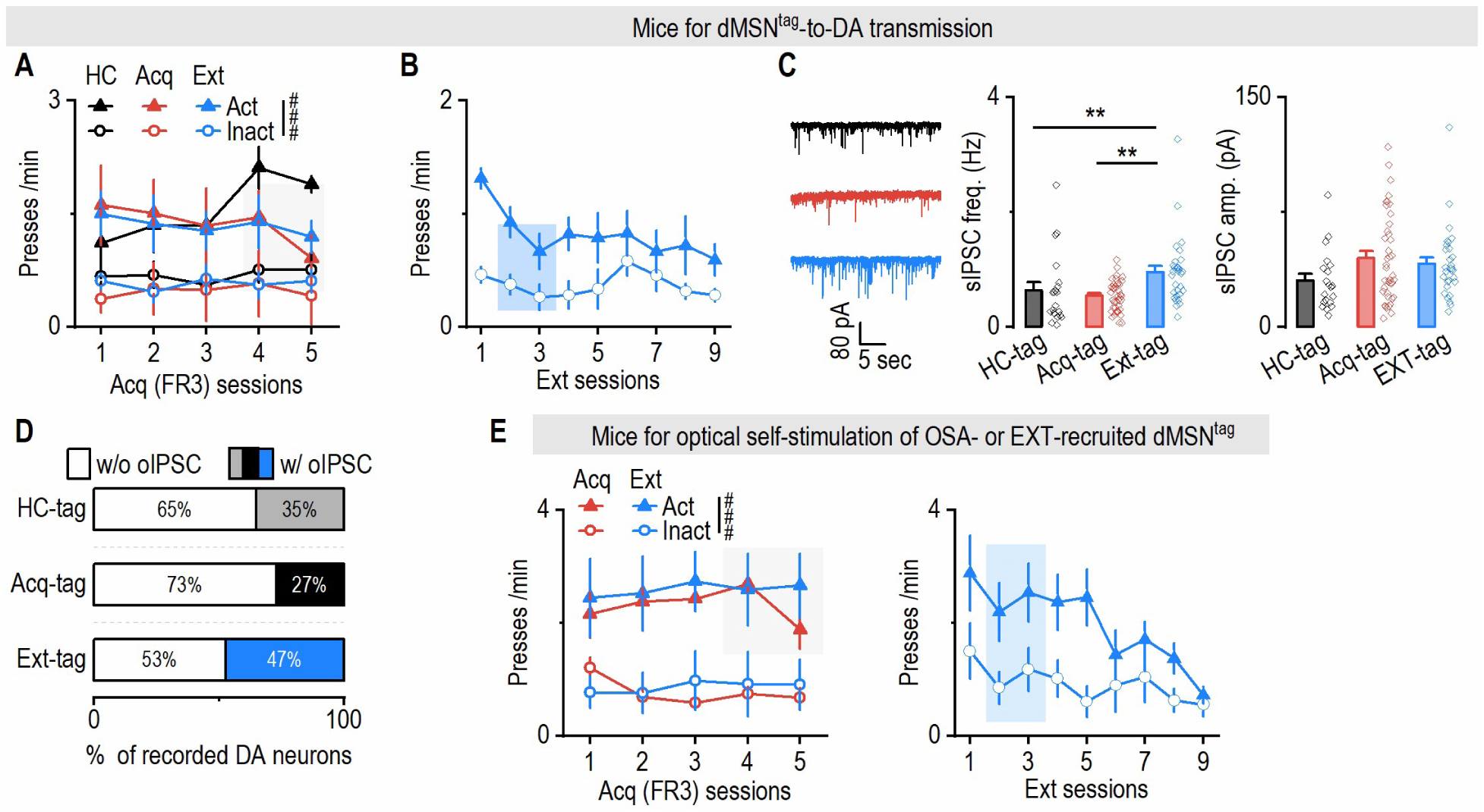
Behavioral performance of mice used in Figure 2. ***A***, Active presses were similar among the three groups (Two-way RM ANOVA: Group: *F*_(2,10)_ = 0.12, *p* = 0.89), and were higher than their inactive presses (Two-way RM ANOVA: Lever: *F*_(1,12)_ = 46.78, ^###^*p* < 0.001). ***B***, In the Ext group, active presses gradually decreased to the level of inactive presses over the course. Two-way RM ANOVA: Lever × Session: *F*_(8,32)_ = 1.76, *p* = 0.12. Sidak’s multiple comparisons test: For sessions 1-9: *t* = 5.84, 3.78, 2.76, 3.69, 3.10, 1.69, 1.46, 2.78, 2.14, *p* < 0.001 for session 1, *p* < 0.01 for sessions 2 and 4, *p* < 0.05 for session 5, *p* > 0.05 for sessions 3 and 6-9. ***C***, SNc DA neurons exhibited similar sIPSC amplitudes across groups (Kruskal-Wallis test: *H*_(2)_ = 4.73, *p* = 0.09). ***D***, A greater percentage of recorded SNc DA neurons received oIPSCs from dMSN^tag^ in the Ext-tag group to the HC-tag and Acq-tag groups. Chi-square test: *Χ*^2^_(2)_ = 3.64, *p* = 0.16. ***E***, For mice tested for oICSS, active presses during the acquisition phase (left) were similar between groups (Three-way ANOVA: Group: *F*_(1,8)_ = 0.17, *p* = 0.69), and exceeded the level of inactive presses (Three-way ANOVA: Lever: *F*_(1,8)_ = 29.03, ^###^*p* < 0.001). During extinction (middle), active presses progressively declined to the level of inactive presses over the course (Two-way RM ANOVA: Lever × Session: *F*_(8,32)_ = 4.05, *p* = 0.002. Sidak’s multiple comparisons test: For sessions 1-9: *t* = 5.19, 5.04, 5.12, 5.08, 6.94, 2.03, 2.51, 2.81, 0.63, *p* < 0.001 for sessions 1-5, *p* > 0.05 for sessions 6-9). The inactive presses (right) during the last acquisition session (Acq-tag) and the last extinction session (Ext-tag) were comparable (Unpaired t test: *t*_(7)_ = 0.102, *p* = 0.92). n = 3 mice (A, B; HC-tag), 5 mice per group (A, B; Acq-tag and Ext-tag), 20 neurons from 3 mice (20/3; HC-tag; C), 40/5 (Acq-tag; C), 30/5 (Ext-tag; C), 20/3 (HC-tag; D), 48/5 (Acq-tag; D), 36/5 (Ext-tag; D), 4 mice (Acq-tag; E) and 5 mice (Ext-tag; E).

**Supplementary Figure 3.**
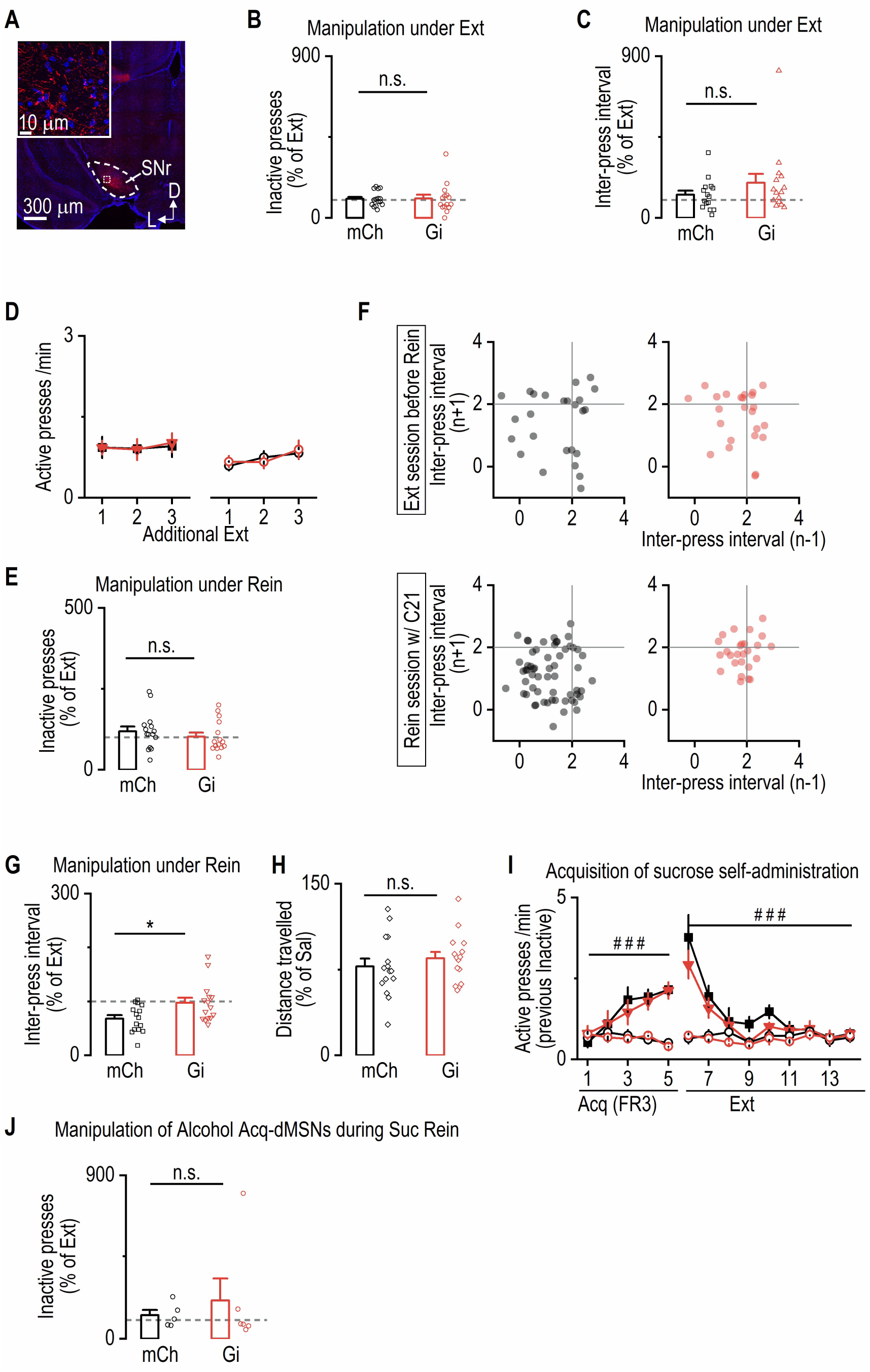
Other behavioral parameters during chemogenetic inhibition of alcohol acquisition-recruited dMSNs (Related to Figure 3) ***A*,** Sample confocal images showing hM4Di-mCherry^+^ fibers, but not cell bodies, in the SNr. D, dorsal; L, lateral. ***B*,** Chemogenetic inhibition of Acq-recruited dMSNs during extinction did not alter inactive-pressing. Mann-Whitney test: *U* = 101, *p* = 0.653. ***C*,** Chemogenetic inhibition of alcohol acquisition-recruited dMSNs during the extinction test did not alter the inter-press interval for active lever between groups. Mann-Whitney test: *U* = 84, *p* = 0.25. ***D*,** Both groups exhibited similar levels of active and inactive presses during additional extinction sessions. Three-way ANOVA: Group: *F*_(1,28)_ = 0.01, *p* = 0.91. ***E*,** Chemogenetic inhibition of Acq-recruited dMSNs during cue-induced reinstatement did not alter inactive pressing compared to the mCh group. Mann-Whitney test: *U* = 94, *p* = 0.455. *F, G*, Sample inter-press intervals analyzed from the last additional extinction session (F, top) and cue-induced reinstatement (F, bottom) from one mouse in each group. Chemogenetic inhibition of acquisition-recruited dMSNs increased inter-press intervals compared to the mCh group (G: unpaired t test: *t*_(28)_ = 2.56, \**p* = 0.016). ***H*,** Chemogenetic manipulation of alcohol-recruited dMSNs did not affect the distance traveled in an open field test. Unpaired t test: *t*_(28)_ = 0.79, *p* = 0.43. ***I*,** Both groups showed similar increases in active presses during 5 d of sucrose training (Three-way ANOVA: Lever × Session: *F*_(4,36)_ = 16.06, ^###^*p* < 0.001), followed by significant reductions during extinction (Three-way ANOVA: Lever × Session: *F*_(8,72)_ = 31.91, ^###^*p* < 0.001). ***J*,** Chemogenetic inhibition of alcohol acquisition-recruited dMSNs did not affect inactive presses during cue-induced reinstatement of sucrose seeking. Mann-Whiteney test: *U* = 13, *p* = 0.79. n = 15 mice per group (B-E, G, H), 5 mice (I, J; mCh), 6 mice (I, J; Gi).

**Supplementary Figure 4.**
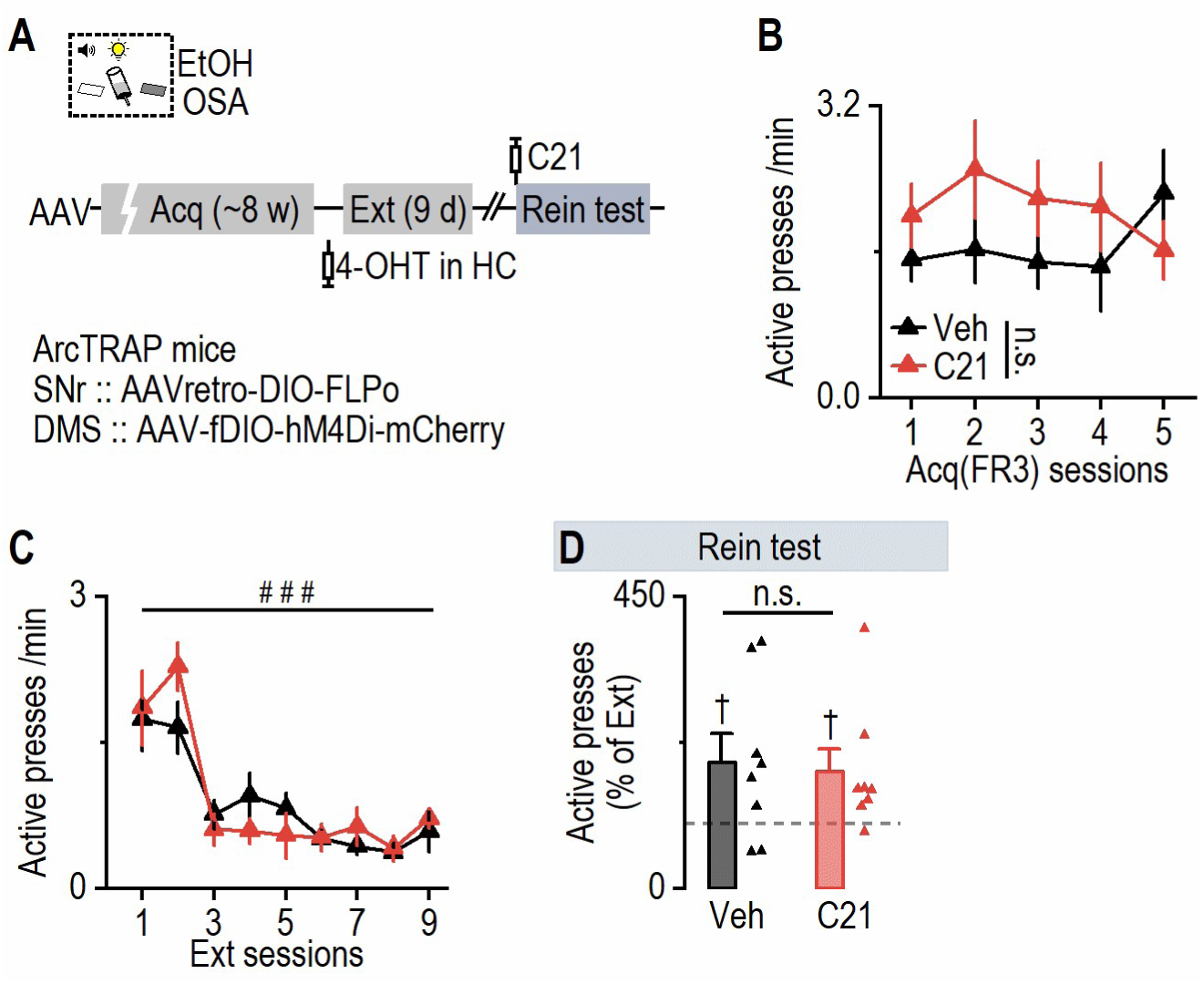
Chemogenetic inhibition of home-cage-recruited dMSNs did not mediate a cue-alcohol memory retrieval (Related to Figure 3) ***A*,** Schematic of the experiment timeline. ArcTRAP mice were infused with AAVretro-DIO-FLPo into the SNr and AAV-fDIO-hM4Di-mCherry into the DMS. Mice were trained for EtOH OSA for ∼8 weeks. At least 24 h after the last OSA session, mice were administered 4-OHT in the home-cage for two consecutive days. One week later, mice underwent 9 d of extinction training, followed by a cue-induced reinstatement test with C21 pre-injection. ***B*,** Active presses were similar between groups. Two-way RM ANOVA: Group: *F*_(1,14)_ = 0.74, *p* = 0.40. ***C*,** Extinction training significantly reduced active presses in both groups. Two-way RM ANOVA: Session: *F*_(8,112)_ = 20.1, ^###^*p* < 0.001. ***D*,** Chemogenetic inhibition of home-cage-recruited dMSNs did not alter cue-induced reinstatement of alcohol seeking. Compared to 100% theoretical baseline: one-sample t test: Veh: *t*_(7)_ = 3.06, ^†^*p* = 0.01; C21: *t*_(7)_ = 2.32, ^†^*p* = 0.053. Group comparison: Mann-Whitney test: *U* = 23, *p* = 0.38. n = 8 mice per group.

**Supplementary Figure 5.**
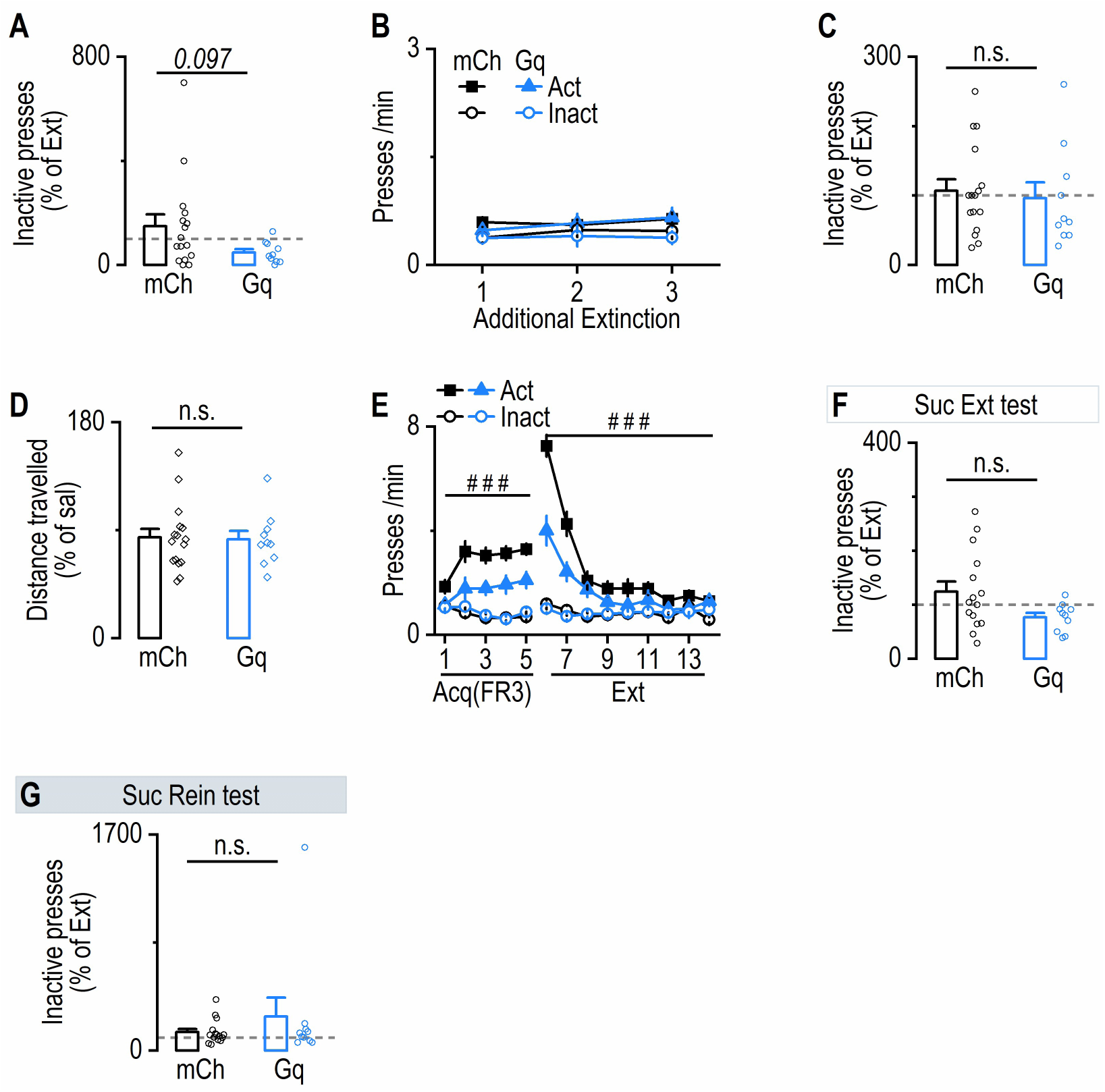
Additional behavior and locomotor activity of mice used in Figure 4) ***A***, During the extinction test, activation of extinction-recruited dMSNs marginally reduced normalized inactive presses compared to mCh. Mann-Whitney test: *U* = 48, *p* = 0.097. ***B***, Both groups showed similar active and inactive presses during 3 d of additional extinction training. Three-way ANOVA: Group: *F*_(1,24)_ = 0.18, *p* = 0.68. ***C***, Activation of extinction-recruited dMSNs did not affect inactive presses during the reinstatement test. Mann-Whitney test: *U* = 67.5, *p* = 0.52. ***D***, Chemogenetic activation of extinction-recruited dMSNs did not alter locomotor activity compared to the mCh group. Mann-Whitney test: *U* = 79, *p* = 0.979. ***E***, Both groups successfully acquired sucrose self-administration, as indicated by significantly higher active versus inactive lever presses across the 5-day training period (Mixed-effect analysis: Session × Lever: *F*_(4,90)_ = 15.77, ^###^*p* < 0.001. Session × Group: *F*_(4,92)_ = 0.28, *p* = 0.89). During extinction training, active lever presses declined in both groups (three-way ANOVA: Session × Lever: *F*_(8,184)_ = 59.06, ^###^*p* < 0.001). ***F***, Chemogenetic activation of EtOH extinction-recruited dMSNs did not alter inactive presses during sucrose extinction test. Unpaired t test: *t*_(23)_ = 1.97, *p* = 0.06. ***G***, Chemogenetic activation of EtOH extinction-recruited dMSNs did not alter inactive presses during sucrose-cue-induced reinstatement test. Mann-Whitny test: *U* = 72, *p* = 0.89. n = 16 mice (mCh; A-D), 10 mice (Gq; A-D), 15 mice (mCh; E-F) and 10 mice (Gq; E-F).

**Supplementary Figure 6.**
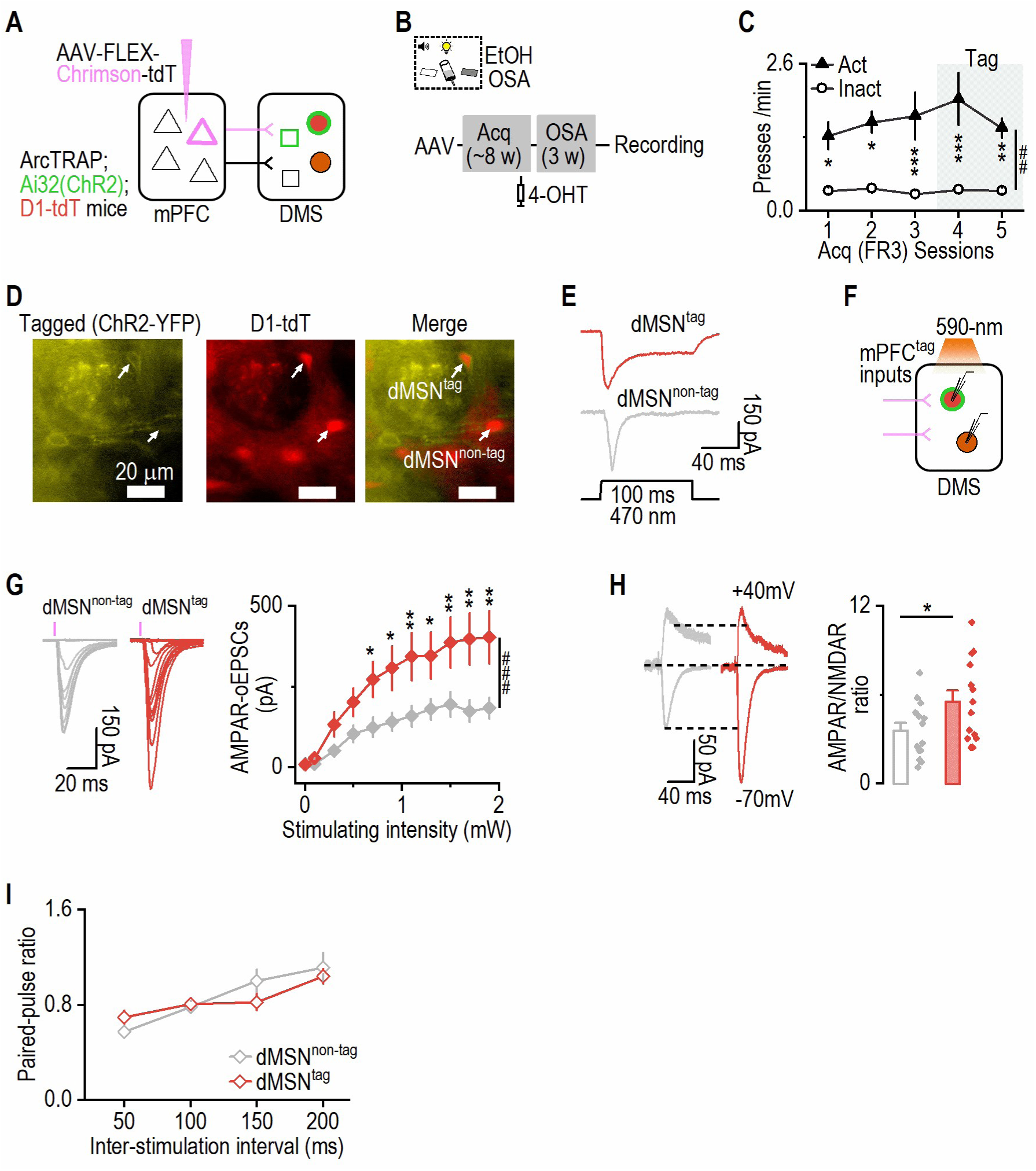
mPFC^tag^-to-dMSN^tag^ transmission, and paired-pulse ratio in ArcTRAP;Ai32;D1-tdT mice (Related to Figure 5) ***A*,** Schematic showing ChR2-EYFP and Chrimson-tdT expression in tagged corticostriatal synapses of ArcTRAP;Ai32 (ChR2-EYFP);D1-tdT mice. AAV-FLEX-Chrimson-tdT was infused into the mPFC to measure optically-induced excitatory postsynaptic currents (oEPSCs) from tagged mPFC neurons (mPFC^tag^) to tagged dMSNs (dMSN^tag^) versus non-tagged dMSNs (dMSN^non-tag^). In the DMS, circle represents dMSN and square represents non-dMSN. ***B*,** EtOH acquisition-activated neurons were tagged during two FR3 sessions, training continued for 3 weeks, and mice were sacrificed 1 d after the last training for electrophysiology. ***C*,** Comparison of active and inactive presses during training and tagging sessions. Mixed-effect analysis: Act vs. Inact: *F*_(1, 12)_ = 16.73, ^##^*p* = 0.0015. Sessions 1-5: *t* = 2.86, 3.05, 4.07, 4.72, 3.26, \**p* < 0.05 (1, 2), \*\*\**p* < 0.001 (3, 4), and \*\**p* < 0.01 (5). ***D*,** Images acquired during electrophysiology recording showing a visually identified dMSN^tag^ (co-expressing ChR2-EYFP and tdT) and a nearby dMSN^non-tag^ (tdT only). ***E*,** Sample traces demonstrate direct depolarization in the ChR2-expressing dMSN^tag^, while the dMSN^non-tag^ shows only oEPSCs in response to 100 ms blue light stimulation. ***F*,** Schematic of synaptic transmission recording from mPFC^tag^ to DMS dMSN^tag^ or dMSN^non-tag^ under 590 nm light stimulation. ***G*,** AMPA receptor-mediated oEPSCs (AMPAR-oEPSCs) from acquisition-activated mPFC inputs were stronger in dMSN^tag^ than dMSN^non-tag^. Mixed-effect analysis: Group × Intensity: *F*_(10, 340)_ = 5.23, ^###^*p* < 0.001; \**p* < 0.05, \*\**p* < 0.01 as compared to dMSN^non-tag^group under the same stimulation intensity. ***H*,** AMPAR/NMDAR ratios (measured with oEPSCs from acquisition-activated mPFC inputs) were greater in dMSN^tag^ compared to dMSN^non-tag^. Unpaired t test: *t*_(26)_ = 2.19, \**p* = 0.038. ***I*,** The paired-pulse ratio (measured by stimulating EtOH acquisition-activated mPFC inputs) did not differ between dMSN^non-tag^ and dMSN^tag^. Two-way RM ANOVA: Group: *F*_(1, 34)_ *=* 0.16, *p* = 0.69. n = 7 mice (C), 19 neurons from 7 mice (19/7; dMSN^non-tag^; G), 17/7 (dMSN^tag^; G), 14/7 per group (H), 18/7 per group (I).

**Supplementary Figure 7.**
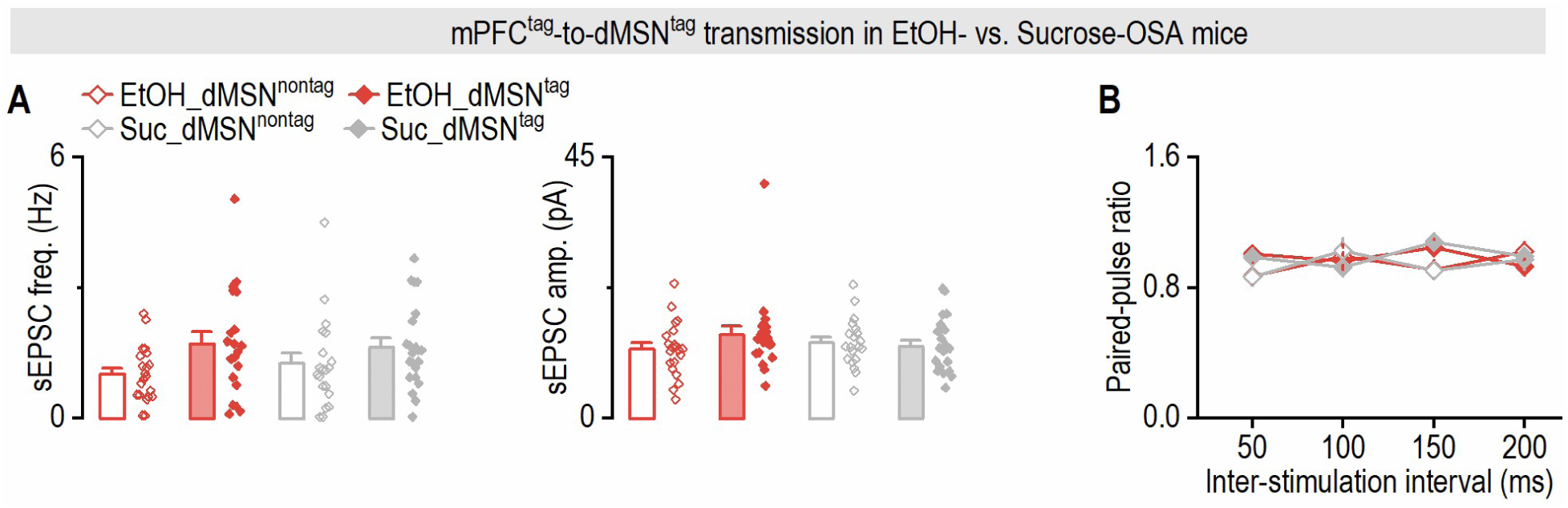
Spontaneous excitatory synaptic transmission and paired-pulse ratio in alcohol- versus sucrose-trained mice (Related to Figure 5) ***A*,** The frequency (left; two-way ANOVA: *F*_(1,79)_ = 0.54, *p* = 0.466) and amplitude (right; two-way ANOVA: *F*_(1,79)_ = 1.79, *p* = 0.18) of sEPSCs did not differ between dMSN^non-tag^and dMSN^tag^ in either the EtOH or sucrose groups. ***B*,** The paired-pulse ratio (measured by stimulating EtOH or sucrose acquisition-activated mPFC inputs) did not differ between dMSN^non-tag^ and dMSN^tag^. Mixed-effect analysis: Substance × Tag: *F*_(1, 107)_ *=* 0.058, *p* = 0.81. n = 21 neurons from 5 mice (21/5; EtOH_dMSN^non-tag^; A), 20/5 (EtOH_dMSN^tag^; A), 20/5 (Suc_dMSN^non-tag^; A), 22/5 (Suc_dMSN^tag^; A), 20/5 (EtOH_dMSN^non-tag^, EtOH_dMSN^tag^, Suc_dMSN^non-tag^; B), and 22/5 (Suc_dMSN^tag^; B).

**Supplementary Figure 8.**
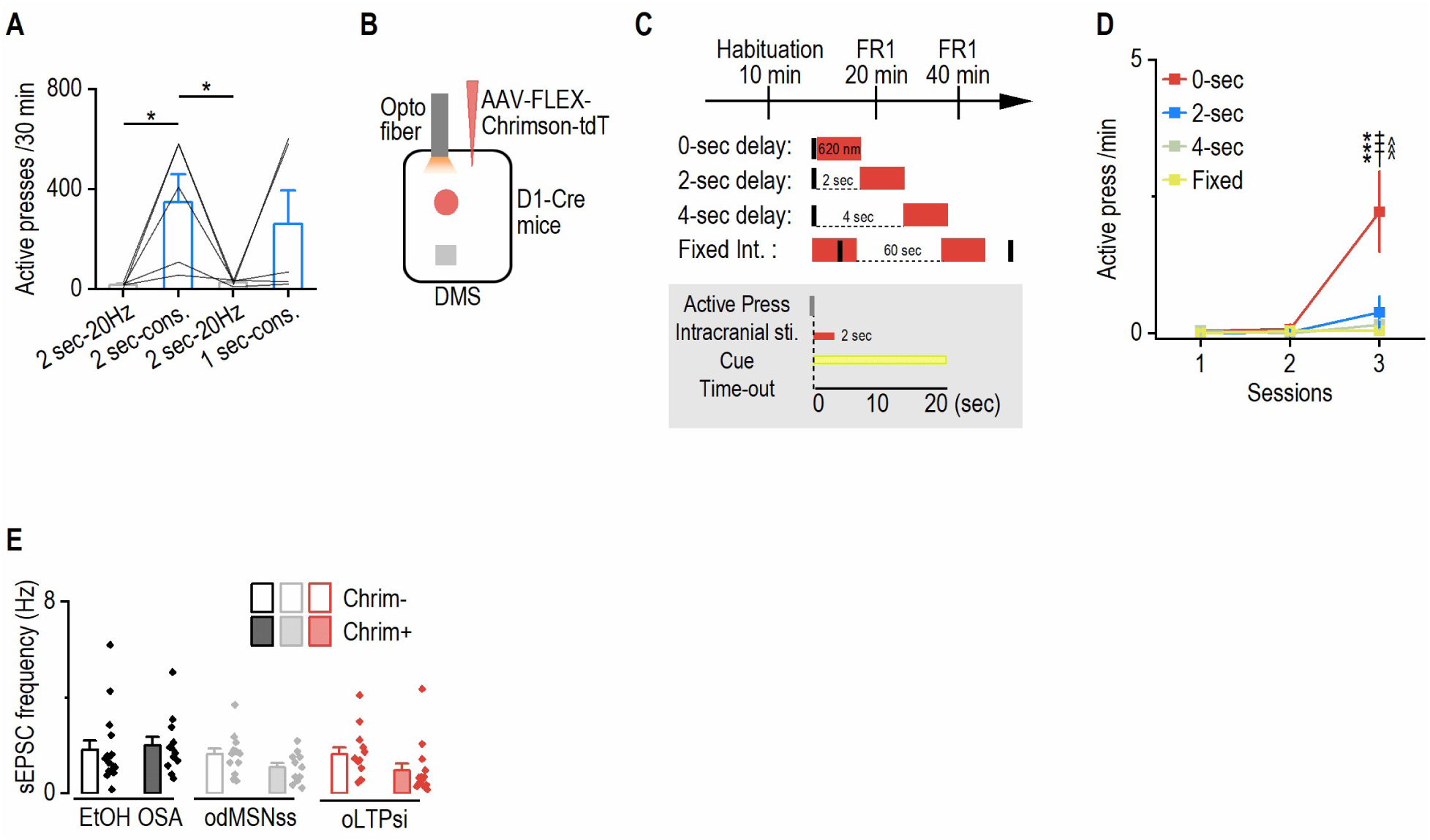
The effect of different odMSNss protocols on operant lever-pressing (Related to Figure 6) ***A***, Control experiments were performed to assess potential optical bleed-through between opsins. Group 1: Wild-type mice received AAV-Chronos-GFP infusions into the mPFC with optic fibers implanted above the DMS. During training, these mice received the postsynaptic stimulation protocol using 590-nm constant light, the wavelength used in Figure 6 to depolarize dMSNs, which should not activate Chronos. Group 2: D1-Cre mice received AAV-FLEX-Chrimson-tdT infusions into the DMS with optic fibers implanted above the DMS. During training, these mice received the presynaptic stimulation protocol using 473-nm, 50-Hz light, the wavelength used in Figure 6 to activate Chronos-expressing mPFC terminals, which may activate Chrimson but to a much less extent. Both groups underwent one day of mock induction (habituation; no light) followed by 3 d of FR1 self-stimulation training, in which each active press triggered the same light-cue stimulation protocol used in Figure 6. ***B***, Neither inappropriate wavelength produced self-stimulation behavior. Specifically, 590-nm light failed to drive self-stimulation in Chronos-expressing mice, and 473-nm light failed to drive self-stimulation in Chrimson-expressing mice, demonstrating minimal functional cross-activation under our experimental conditions. Three-way ANOVA: Lever × Session: *F*_(3, 24)_ = 0.55, *p* = 0.65. n = 5 mice per group. ***C***, Three different stimulation protocols for odMSNss were tested: 2-sec 20-Hz pulsed light, 2-sec constant light, and 1-sec constant light. Only 2-sec constant light stimulation caused an increase in active presses. One-way RM ANOVA: *F*_(3, 12)_ = 4.99, *p* = 0.018. Tukey’s multiple comparisons test: 2-sec 20 Hz vs. 2-sec constant: *q* = 4.42, \**p* = 0.038; 2-sec constant vs. 2-sec 20 Hz again: *q* = 4.31, \**p* = 0.044. n = 4 rats. ***D***, Schematic illustrating AAV-FLEX-Chrimson-tdT infusion into the DMS of D1-Cre mice with bilateral fiber implantation above the infusion sites. ***E***, After one day of habituation, different stimulation protocols for odMSNss were tested across 2 d of FR1: 0-sec delay (light stimulation immediately following an active press), 2-sec delay (2-sec delay between the active press and light stimulation), 4-sec delay, and fixed interval stimulation (light delivered every 60 sec, regardless of active presses). ***F***, Only the 0-sec delay group exhibited self-stimulation after 2 d of training. Two-way RM ANOVA: Protocol × Session: *F*_(6,12)_ = 4.21, *p* = 0.016. Tukey’s multiple comparisons test: In session 3: 0-sec vs. 2-sec: *q* = 6.93, \*\*\**p* < 0.001; 0-sec vs. 4-sec: *q* = 6.94, ^†††^*p* < 0.001; 0-sec vs. fixed interval: *q* = 7.41, ^^^^^*p* < 0.001. n = 3 mice per group (0-sec and 2-sec), 2 mice per group (4-sec and fixed interval). ***G***, sEPSC frequencies did not differ between Chrim^−^ and Chrim^+^ neurons across all three groups. Two-way ANOVA: Group × Neuron (Chrim^-^ vs. Chrim^+^): *F*_(2,73)_ = 2.47, *p* = 0.32. n = 16 neurons/4 mice (Chrim^-^ in EtOH), 12/4 (Chrim^+^ in EtOH), 13/3 (Chrim^-^ in odMSNss), 11/3 (Chrim^+^ in odMSNss), 13/3 (Chrim^-^ in oLTPsi), 14/3 (Chrim^+^ in oLTPsi).

**Supplementary Figure 9.**
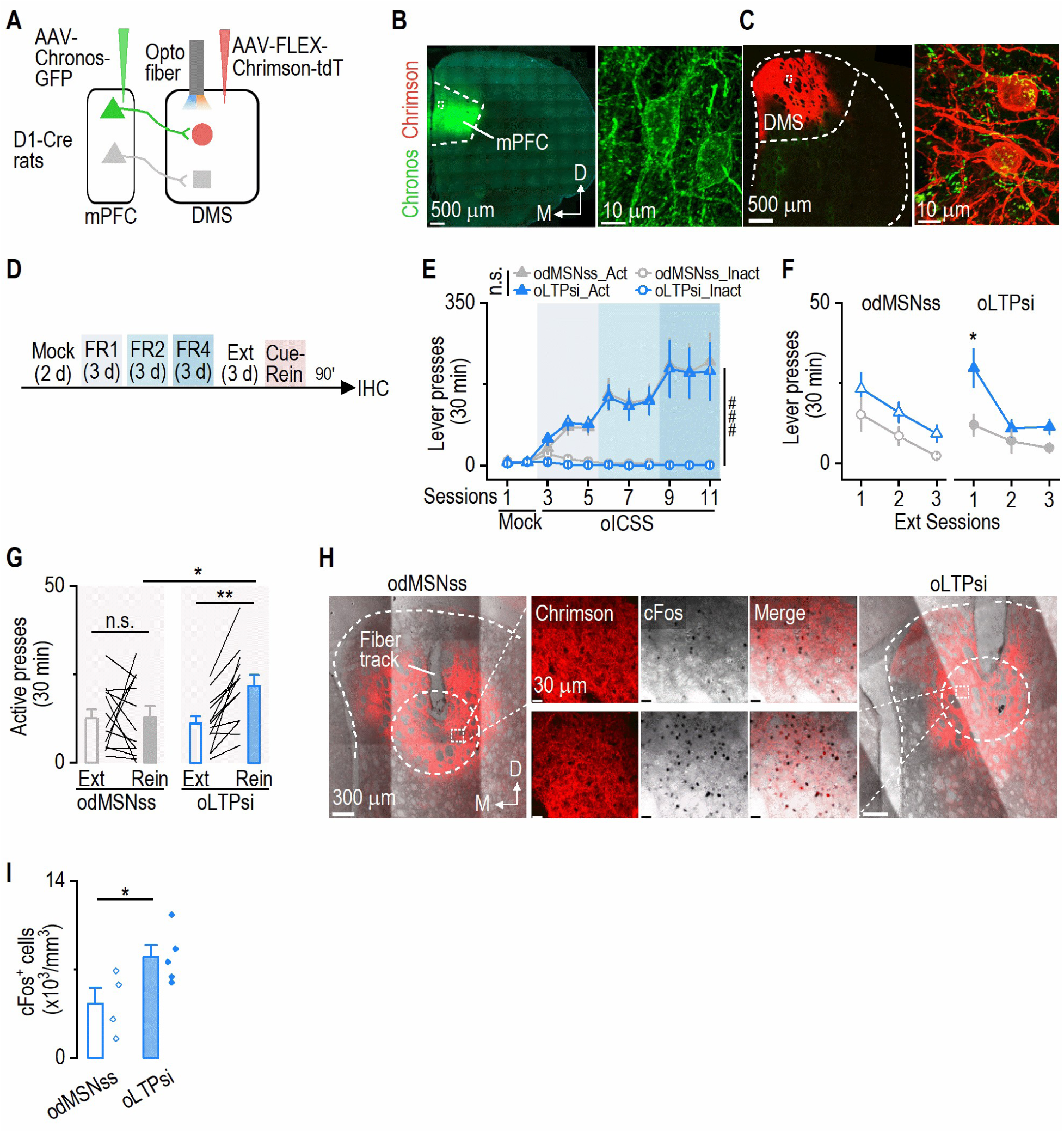
oLTPsi and odMSNss in D1-Cre rats (Related to Figure 6) ***A***, The schematic of virus infusion and fiber implantation in D1-Cre rats. ***B, C***, Sample images showing Chronos-GFP expression in the mPFC (B) and Chrimson-tdT expression in the DMS (C). Enlarged images (right) showed individual neurons from the outlined regions (left). D, dorsal; M, medial. ***D***, Timeline for optogenetic intracranial self-stimulation (oICSS) using either oLTPsi or odMSNss. ***E***, Active and inactive presses during the training period were similar between odMSNss and oLTPsi groups (Three-way ANOVA: Group: *F*_(1,48)_ = 0.03, *p* = 0.86). Both groups performed more active presses than inactive presses (Three-way ANOVA: Lever: *F*_(1,48)_ = 34.34, ^###^*p* < 0.001). ***F***, The oLTPsi group, but not the odMSNss group, continued to display more active than inactive presses during the first extinction session. Three-way ANOVA: Group × Lever × Session: *F*_(2,96)_ = 1.65, *p* = 0.19. Tukey’s multiple comparisons test: Act vs. Inact during first extinction in oLTPsi: *q* = 4.98, \**p* = 0.027; Act vs. Inact during first extinction in odMSNss: *q* = 2.23, *p* = 0.91. ***G***, The oLTPsi group, but not the odMSNss group, showed cue-induced reinstatement of active pressing. Two-way RM ANOVA: Group × Session: *F*_(1,24)_ = 6.07, *p* = 0.021. Sidak’s multiple comparisons test: Rein vs. Ext: *t* = 0.10, *p* = 0.99 (odMSNss); *t* = 3.59, \*\**p* = 0.003 (oLTPsi). odMSNss vs. oLTPsi in Rein: *t* = 2.37, \**p* = 0.043. ***H***, Sample confocal images of striatal sections from an odMSNss and an oLTPsi rat containing c-Fos^+^ neurons in the Chrimson-expressing region below the fiber track. Dashed circles indicate regions of analysis and dashed rectangles indicate where the enlarged images were taken from. ***I***, The oLTPsi group had more c-Fos^+^ neurons than the odMSNss group in the DMS regions with high Chrimson-tdT expression. *t*_(7)_ = 2.40, \**p* = 0.047. n = 13 rats/group (E-G), 4 rats (I; odMSNss), and 5 rats (I; oLTPsi).

## Materials and Methods

### Reagents

AAVretro-pEF1a-DIO-FLPo (#87306), AAV8-hSyn-fDIO-hM4Di-mCherry (#154867), AAV8-hSyn-fDIO-hM3Dq-mCherry (#154868), AAV8-Ef1a-fDIO-mCherry (114471), AAV8-Ef1a-DIO-BFP2.0 (#137130) were obtained from Addgene. AAV8-Syn-Chronos-GFP, AAV8-Syn-FLEX-Chronos-GFP, AAV8-Syn-FLEX-Chrimson-tdTomato, and AAV5-Ef1a-DIO-hsChRmine-oScarlet were purchased from the University of North Carolina Vector Core. NBQX and APV were purchased from R&D Systems. Picrotoxin, 4-hydroxytamoxifen (4-OHT), DREADD agonist compound 21 (C21), and other reagents were either purchased from Sigma, or gifted by the National Institute on Mental Health Drug Supply Program (4-OHT and C21).

### Animals

Drd1a-Cre (D1-Cre) mice were obtained from the Mutant Mouse Regional Resource Centers ^24,74^, Drd1a-tdTomato ^75^, Ai140-GFP ^57,76,77^, Snap25-GFP ^78^, ArcTRAP (CreER^T2^) ^27,39,76^ mice were purchased from the Jackson Laboratory. D1-Cre rats were obtained from the Rat Resource and Research Center^79^. ArcTRAP mice were crossed with Ai140-GFP and D1-tdTomato mice to obtain ArcTRAP;Ai140-GFP;D1-tdTomato mice. Genotypes were determined by PCR analysis of Cre or the fluorescent protein gene in tail DNA (Cre for D1-Cre mice; tdTomato for D1-tdTomato mice; GFP for Ai140-GFP^80,81^. All animals that were used are in mixed gender and aged from 3-8 months. Animals were housed in a temperature- and humidity-controlled vivarium with a reversed 12-h light/dark cycle (lights on at 10:00 P.M. and off at 10:00 A.M.). All behavior experiments were conducted in their dark cycle, starting approximately 1 h after the light went off. Food and water were available *ad libitum*. All animal care and experimental procedures were approved by the Texas A&M University Institutional Animal Care and Use Committee.

## Experiment procedures

### Intermittent access to 20% alcohol two-bottle-choice drinking procedure

To establish high levels of alcohol consumption in mice, we utilized the intermittent-access to 20% alcohol two-bottle-choice drinking procedure ^24,57,68,80,82^. ArcTRAP and ArcTRAP;Ai140-GFP;D1-tdTomato mice were given free access to two bottles containing water or 20% alcohol for three 24-h sessions (Mondays, Wednesdays, and Fridays), with 24-h or 48-h withdrawal periods (Tuesdays, Thursdays, Saturdays, and Sundays) each week. During the withdrawal periods, the mice had unlimited access to water bottles. The placement of the alcohol bottle was alternated for each drinking session to control side preferences. For all animals that underwent operant self-administration of alcohol, 3-6 weeks of 2BC were used to acclimate them with alcohol.

### Operant self-administration of alcohol

Animals were trained to self-administer 20% alcohol in operant chambers using a fixed ratio (FR) schedule ^43,56,83^. The mouse operant setup (Med Associates) included two levers in each chamber: active and inactive, on the opposite sides of the same wall.

During alcohol OSA training sessions, a house light centered above the operant chamber was illuminated. Each operant experiment commenced with continuous reinforcement (FR1) every other day for approximately two to three weeks, wherein pressing the active lever resulted in a ∼0.32 mL delivery of 20% alcohol. Actions on the inactive side were documented but did not initiate a programmed event. The alcohol solution was dispensed into a stainless-steel reservoir situated within the magazine port between active and inactive levers. Each alcohol delivery persisted for 3 sec and was accompanied by a tone and a discrete yellow cue light in the magazine port for the same duration. There was a 20-sec time-off period for each delivery, during which active/inactive presses were recorded but did not cause any further alcohol deliveries. When animals were able to achieve at least 10 deliveries under the FR1 schedule, the criterion to receive the alcohol escalated to FR2 (around one to two weeks), and eventually progressed to FR3 (around two weeks). Each operant session for mice lasted 30 min or 1 h, depending on the learning condition.

### Extinction training

The extinction training was conducted under the FR3 schedule. However, pressing on the active side did not result in actual alcohol delivery, nor cue presentation. Each extinction session for mice lasted 30 min, and in total lasted for 9 d. For chemogenetic manipulations during the extinction, C21 (i.p., 1 mg/kg; or saline) was injected 30 min prior to the start of the test session.

### Cue-induced reinstatement of alcohol seeking

The reinstatement test (30 min) was conducted as previously described^67^. It was carried out at the FR3 schedule, where three actions on the active side triggered the illumination of the cue light within the magazine port, the tone, and the pump action sound, with the exception that the first cue presentation was given immediately after the first active press. For chemogenetic manipulations during reinstatement, C21 (i.p., 1 mg/kg; or saline) was injected 30 min prior to the start of the test session.

### Operant self-administration of sucrose

For chemogenetic manipulation of alcohol-recruited dMSNs during sucrose reinstatement, mice were trained directly under the FR3 schedule. For each animal, the previous inactive lever for alcohol was assigned as the active lever for sucrose, and vice versa. Sucrose pellets (20 mg/pellet/delivery, Bio-Serv #F07595) were delivered into the stainless-steel cup next to the alcohol port. To differentiate the alcohol cue with the sucrose cue, each sucrose delivery was accompanied by distinct green cue light with no tone delivery. Each delivery also had a 20-sec time-out period, during which active presses did not cause any further deliveries.

For electrophysiological recording of mPFC^tag^-to-dMSN^tag^ or -dMSN^non-tag^ transmission, mice were trained for either alcohol or sucrose OSA under the FR1 schedule for 10 d. Each alcohol/sucrose delivery persisted for 3 sec, accompanied by a tone and a discrete yellow cue light in the magazine port for the same duration. There was a 20-sec time-off period for each delivery, during which active/inactive presses were recorded but did not cause any further alcohol deliveries. After the lasting training session, mice were returned to their home-cages for 21 d until recording.

### Tag active neurons

4-OHT was prepared and tagging was conducted as previously described ^27,76,84^. The 4-OHT solution was freshly prepared on the tagging day. In brief, 4-OHT powder was dissolved in 200 proof ethanol (20 mg/mL) at room temperature, with continuous shaking until the 4-OHT was fully dissolved. An equal volume of a 1:4 mixture of castor oil and sunflower seed oil was added, resulting in a final concentration of 10 mg/mL 4-OHT.The ethanol was removed by centrifugation under vacuum. The final 4-OHT in oil was stored at 4°C for a maximum of 12 h before use.

For tagging during acquisition or extinction of operant learning, mice were habituated to i.p. injection of saline immediately after each operant session for two consecutive days. On the third and fourth operant sessions (Tag), mice were introduced to an isoflurane chamber immediately after the operant session and 4-OHT was i.p. injected. The use of isoflurane is to reduce the pain and discomfort due to 4-OHT/oil mix injection. After the injection, mice were returned to their home-cage and left without any disturbance for the following 6 hours. For tagging in the home-cage condition, mice were habituated to i.p. saline injections for two consecutive days, beginning ∼24 hours after the previous operant training. During tagging days (first tagging day was ∼24 h after the last operant session), mice received an i.p. injection of 4-OHT while remaining in their home-cages. A second 4-OHT injection was administered 24 hours later.

ArcTRAP was used for activity-dependent tagging because it robustly labels learning-activated neurons in the dorsal striatum, whereas FosTRAP labels relatively few striatal neurons^39^; neuronal reactivation was assessed using c-Fos immunohistochemistry, which provides a reliable and widely validated marker of recent neuronal activity.

### Stereotaxic virus infusion and fiber implantation

Where required for the experimental design, AAVretro-DIO-FLPo (0.6 µL/site) was bilaterally infused into the SNr (AP: -3.15 mm; ML: ±1.35 mm; DV: -4.6 mm), AAV-Chronos-GFP was infused into the mPFC (AP: 1.94 mm; ML: ± 0.3mm; DV: -2.5 mm), and/or AAV-FLEX-Chrimon-tdT (1 µL/site) was bilaterally infused into the DMS (AP: +0.3 mm; ML: ±1.87 mm; DV: -2.9 to -3.1 mm) of D1-Cre mice, and AAV-fDIO-mCherry/AAV-fDIO-hM3Dq-mCherry/AAV-fDIO-hM4Di-mCherry/AAV-DIO-BFP/AAV-DIO-ChRmine-oScarlet (0.5 µL/site) was bilaterally infused into the DMS (AP1: +0.98 mm; ML1: ±1.25 mm; DV1: -2.9 to -3.1 mm; AP2: +0.3 mm; ML2: ±1.87 mm; DV2: -2.9 to -3.1 mm) of ArcTRAP mice. Optical fibers implants (200-μm diameter optical fiber secured to a 1.25-mm stainless steel ferrule) were placed bilaterally in DMS through the injection tract (DV: -2.85 mm) or in the SNc/SNr area (AP: -3.15 mm; ML: 1.35 mm; DV: -4.55 mm). In rats, AAV-Chronos-GFP (0.8 µL/site) was infused into the mPFC (AP: +2.9 mm, ML: ±0.65 mm, DV: -3.5 mm) and AAV-FLEX-Chrimson-tdT (1.2 µL/site) was infused into the DMS (AP: +0.36 mm, ML: ±2.3 mm, DV: -4.7 mm). Optical fiber implants (300-μm diameter optical fiber secured to a 2.5-mm stainless steel ferrule) were placed bilaterally in the DMS through the injection tract (DV: -4.6 mm). Animals were anesthetized with 3-4% isoflurane at 1.0 L/min and mounted in a stereotaxic surgery frame. The head was leveled and craniotomy was performed using stereotaxic coordinates adapted from the brain atlas ^85^. The virus was infused at a rate of 0.08 µL/min. At the end of the infusion, the injectors remained at the injection site for an additional 10-15 min before removal to allow for virus diffusion. For fiber implants, 2 metal screws (mice) or four metal screws (rats) were fixed into the skull to support the implants, which were further secured with dental cement (Henry Schein). The scalp incision was then sutured, and the animals were returned to their home cage for recovery.

### Slice electrophysiology

#### Slice preparation

Slices were prepared and electrophysiological recordings were conducted as has been described before^43,56,82,83,86^. Briefly, coronal sections containing the striatum (250 μm) were cut in an ice-cold cutting solution containing (in mM): 40 NaCl, 148.5 sucrose, 4 KCl, 1.25 NaH_2_PO_4_, 25 NaHCO_3_, 0.5 CaCl_2_, 7 MgCl_2_, 10 glucose, 1 sodium ascorbate, 3 sodium pyruvate, and 3 myo-inositol. The solution was saturated with 95% O_2_ and 5% CO_2_. Slices were then incubated in a 1:1 mixture of the cutting solution and external solution at 32°C for 40-45 min. The external solution was composed of the following (in mM): 125 NaCl, 4.5 KCl, 2.5 CaCl_2_, 1.3 MgCl_2_, 1.25 NaH_2_PO_4_, 25 NaHCO_3_, 15 glucose, and 15 sucrose. The external solution was saturated with 95% O_2_ and 5% CO_2_. Slices were then maintained in the external solution at room temperature until use. Striatal slices were prepared 1-3 d after the last acquisition session in ArcTRAP;Ai32;D1-tdTomato mice, 21 d after the last acquisition session in ArcTRAP;D1-tdTomato mice (for alcohol OSA vs. sucrose OSA), 1-3 d after the last training session in D1-Cre mice trained for alcohol OSA or oICSS, and 2 weeks after the last extinction training for ArcTRAP mice (for dMSN^tag^-DA transmission).

#### Whole-cell recordings

Individual slices were transferred to a recording chamber and continuously perfused with the external solution at a rate of 2-3 mL/min at 32°C. Picrotoxin (100 μM) was included in the external solution of recordings for glutamatergic transmission to block GABA_A_ receptor-mediated transmission. DNQX (20 μM) and D-AP5 (50 μM) were included in the external solution of recordings for GABAergic transmission to block AMPA and NMDA receptor-mediated responses. Fluorescent axonal fibers and neurons were visualized using an epifluorescent microscope (Examiner A1, Zeiss). For measuring synaptic transmission using optogenetic stimulation of presynaptic terminals, we delivered a 2-ms, submaximal (∼1.6 mW) stimulation before recording and excluded neurons that did not show optically induced EPSCs. Neurons were recorded at a uniform depth (∼75 μm) around locations with the strongest virus expression to prevent variability from tissue depth. For ArcTRAP;Ai32;D1-tdTomato mice, prior to patching onto a cell in the DMS, the presence of tdT and/or ChR2-EYFP expression was used to visually verify the cell type, and the expression of ChR2 was functionally verified by delivering a 100 ms 470 nm light stimulation. Once we identified a ChR2-expressing dMSN (dMSN^tag^), a dMSN without ChR2 (dMSN^non-tag^) was also selected as a pair, under the same field of view, in a close-distance (< 100 mm) to the patched dMSN^tag^. For ArcTRAP;D1-tdTomato mice, the presence of tdT and BFP was used to visually identify dMSN^tag^, and a dMSN^non-tag^ was identified under the same field of view in a close-distance (< 100 mm) to the patched dMSN^tag^. For D1-Cre mice, Chrimson^+^ neurons were identified by their tdT fluorescence expression and functionally verified by delivering a 100 ms 590 nm light stimulation. A Chrimson^-^ neuron was then selected under the same field of view, in a close-distance to the patched Chrimson^+^ neuron. For ArcTRAP mice, to measure dMSN^tag^-to-DA transmission, the ventromedial SNc was first located as previous studies found that DMS dMSNs mainly project to this part of SNc. Then, we visually identified the DA neurons based on their morphology (∼20 mm in size). MSNs were clamped at -70 mV and DA neurons were clamped at -60 mV. We used a Cs-based intracellular solution containing (in mM): 119 CsMeSO_4_, 8 tetraethylammonium chloride, 15 4-(2-hydroxyethyl) piperazine-1-ethanesulfonic acid (HEPES), 0.6 ethylene glycol tetraacetic acid (EGTA), 0.3 Na_3_GTP, 4 MgATP, 5 QX-314, and 7 phosphocreatine, with an osmolarity of ∼280 mOsm/L. The pH was adjusted to 7.3 with CsOH.

Once each cell was successfully patched, a 3-min spontaneous EPSC/IPSC was recorded. The last minute of each recording was used to analyze the frequency and amplitude of sEPSCs/sIPSCs. For selective stimulation of inputs from channelrhodopsin-expressing fibers onto DMS neurons, 470-nm or 590-nm light was delivered through the objective lens for 2 ms. To generate input-output curves for AMPAR-mediated oEPSCs or GABAR-mediated oIPSCs, light-evoked currents were recorded at 10 different stimulation intensities. To obtain AMPAR/NMDAR ratios, we initially clamped neurons at -70 mV and measured five EPSCs; neurons were then clamped to +40 mV, and five additional EPSCs were recorded. AMPAR values were calculated by averaging the current traces recorded at -70 mV and measuring the amplitude of the averaged peak. The +40 mV traces were also averaged, and the NMDAR current was determined 30 ms after the peak AMPAR current, when the contribution of the AMPAR component was minimal^56^. The AMPAR/NMDAR ratio was calculated by dividing the peak amplitude of AMPAR-EPSCs by that of NMDAR-EPSCs. To measure the probability of transmitter release, we delivered a pair of light stimulations at an interval of 50, 100, 150, 200 ms. The PPR was calculated by dividing the amplitude of the second EPSC by the amplitude of the first EPSC.

### Optogenetic intracranial self-stimulation

The dMSN self-stimulation was conducted as has been shown before ^25,57,61,63,65^. The designation of the active side was randomized for each animal. During the initial 2-d mock induction period, mice or rats were connected to an optical fiber; however, no stimulation was delivered following each active press. Starting from the FR1 schedule, every active press triggered a delivery of pre- and post-synaptic stimulation pairing for oLTPsi or post-synaptic depolarization alone for odMSNss, accompanied by a 20-sec cue light displayed above the active lever and a tone. The LTP protocol delivers simultaneous pre- and postsynaptic stimulation, in which the cross-activation between blue/yellow light and Chronos/Chrimson does not affect the induction efficacy. A 20-sec time-out period followed each light stimulation. After a 3-d period under the FR1 schedule, animals then underwent training under the FR2 schedule for 3 d and then the FR3 (FR4 for rats) schedule for another 3 d. Upon completion of the FR3 (or FR4) training, some animals were used for electrophysiology recordings while some underwent a 6-d (3-d for rats) extinction training under the FR3 (or FR4) schedule, where active pressing no longer triggered any stimulation or cue. After extinction training, animals were assessed for cue-induced reinstatement. During the test, the cue was delivered under the FR3 (or FR4) schedule, except for the first cue which was non-contingently delivered after the session onset. No stimulation was provided during the reinstatement test.

## Open field locomotor activity test

A transparent open field activity chamber (Med Associates, 43 cm x 43 cm x 21 cm height) was equipped with an infrared beam detector that was connected to a computer. All tests lasted for 10 min. Animals were first habituated in the chamber to remove the novelty-induced effect. Then, animals were tested for locomotor activity by placing them into the chamber 30 min after i.p. injections of saline. One day later, animals were re-introduced to the chamber 30 min after i.p. injections of C21 (1 mg/kg). Locomotor activity during the C21-injected day was compared between groups by normalizing the distance travelled in the C21-injected day to the distance travelled in the saline-injected day.

## Histology and cell counting

Animals were perfused intracardially with 4% formaldehyde in phosphate-buffered saline. The brains were removed and post-fixed overnight in 4% formaldehyde in phosphate-buffered saline at 4°C prior to dehydration in 30% sucrose solution. The brains were cut into 50-µm coronal sections using a cryostat. c-Fos and MOR IHC was carried out on free-floating sections. All incubations (except for the primary antibody) and rinses took place at room temperature on a shaker. Sections were rinsed in PBS three times between steps. Sections were first placed into 0.3% H_2_O_2_ for 15 minutes (only for rat slices), followed by blocking in 2% normal donkey serum in PBS with 0.3% triton-X (PBST) for 1 hour and incubation in primary antibody (rabbit anti-c-Fos: for mice: 1:2000, for rats: 1:10000, EMD-Millipore, cat# PC38; anti-MOR: 1:500, Immunostar #24216) diluted in blocking solution overnight under 4°C. Sections were then incubated in 1:500 biotinylated donkey anti-rabbit (Jackson Immuno Research) for 1 hour. For mice slices, sections were visualized via incubation for 1 hour in Streptavidin-conjugated Alexa 647 (1:1000, Thermo Fisher). For rat slices, sections were incubated in ABC (1:500, Vector Elite Kit, Vector Labs) for 45 minutes and the reaction was visualized via incubation for 10 minutes in 0.025% 3,3’-diaminobenzidine (DAB), 0.05% nickel ammonium sulfate, and 0.015% H_2_O_2_. For anti-MOR, slices were directly incubated in biotinylated donkey anti-rabbit conjugated Alexa 647 (1:500, Thermo Fisher) for 1 hour. After incubation, all slices were washed 3 times in PBS.

All images were acquired using a confocal microscope (FluoView 3000, Olympus, Tokyo, Japan) and analyzed using IMARIS 8.3.1 (Bitplane, Zürich, Switzerland), as previously reported^23,81^. For cell counting in slices from ArcTRAP or FosTRAP mice, analyses were performed within a circular region-of-interest (ROI; radius = 300 µm) positioned consistently at the dorsal-medial boundary of the DMS across all slices. For striosome analysis, MOR^+^ striosomes in the DMS were selected using the contour function in IMARIS and counting was conducted within the striosome. To calculate the estimated dMSN^tag^ density in the matrix, the following formula was utilized based on the published striosome-to-matrix volume ratios.

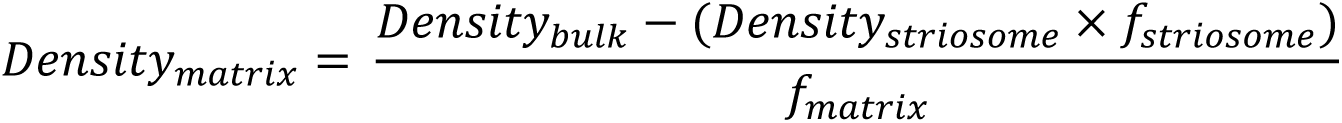

where *f*_striosome_ is 20% and *f*_matrix_ is 80%^35^.

For c-Fos counting in rat slices, analysis was conducted in a circular ROI with a radius of 500 µm directly under the fiber implantation track.

On average, 4-8 slices from the pDMS region (range around AP: 0.62 mm to -0.1 mm from Bregma) were obtained from each mouse; 2-6 slices were obtained from each rat.

## Statistical analysis

All data are presented as mean ± standard error of the mean (SEM). Normality was assessed using the Shapiro-Wilk test. If the data met normality assumptions, statistical comparisons were conducted using a two-tailed t-test (paired or unpaired) or one-, two-, or three-way ANOVA (with repeated measures), followed by post hoc Tukey’s or Sidak’s multiple comparisons test, when appropriate. If normality was not met, data were analyzed using the Mann-Whitney test, Kruskal-Wallis test (followed by Dunn’s post hoc test), or mixed-effects analysis with post hoc Tukey’s or Sidak’s multiple comparisons test. Statistical significance was set at p < 0.05. All statistical analyses were performed using SigmaPlot, and graphs were generated using OriginPro.

## Acknowledgements

We appreciate critical comments on our manuscript from Drs. Stephen Maren, Dorit Ron, Jun Ding, and Yong Xu. We thank the National Institute on Mental Health Drug supply program for giving 4-OHT and C21. This research was supported by NIAAA Grant R01AA021505, R01AA027768, U01AA025932 and X-grant from Texas A&M University to J.W., NIDA Grant R01DA046457 to R.S., McGovern Fellowship from Texas Research Society on Alcoholism (TRSA) to X.X., and Doctoral Student Small Grant from Research Society on Alcohol (RSA) to X.X.

## Author contributions

Conceptualization: J.W., X.X.; Methodology: X.X., J.W., R.S.; Investigation and Formal analysis: X.X., Y.H., R.C., H.G., Z.H., J.L., E.Y., A.C., N.H.; Writing - Original Draft: X.X.; Writing – Review & Editing: J.W., X.X., and V.V; Resources: X.W.; Funding acquisition: J.W., R.S., and X.X.; Supervision: J.W. and R.S.

## Data and materials availability

All data are available in the main text or the supplementary materials.

## Declaration of generative AI and AI-assisted technologies in the writing process

During the preparation of this work, the authors used ChatGPT (OpenAI) to improve the grammar and readability of the manuscript. After using this tool, the authors reviewed and edited the content as needed and took full responsibility for the content of the published article.

## Supplementary Information

### List of acronyms used in the main text (in alphabetical order)

2BC: Two-Bottle Choice
4-OHT: 4-Hydroxytamoxifen
AAV: Adeno-Associated Virus
AMPA: α-amino-3-hydroxy-5-methyl-4-isoxazolepropionic acid
C21: Compound 21 (a DREADD agonist)
DMS: Dorsomedial Striatum
dMSNs: Direct-Pathway Medium Spiny Neurons
DREADDs: Designer Receptors Exclusively Activated by Designer Drugs
EPSCs: Excitatory Postsynaptic Currents
FR: Fixed Ratio
Gi: Inhibitory hM4D (Gi)
Gq: Excitatory hM3D (Gq)
IHC: Immunohistochemistry
IPSCs: Inhibitory Postsynaptic Currents
LTP: Long-Term Potentiation
mCh: mCherry (fluorescent protein control)
MOR: Mu-Opioid Receptor
mPFC: Medial Prefrontal Cortex
NMDA: N-methyl-D-aspartate
oICSS: Optogenetic Intracranial Self-Stimulation
oLTPsi: Self-Induce Optical mPFC-to-dMSN Long-Term Potentiation
odMSNss: Optical dMSN Self-Stimulation
SNc: Substantia Nigra Pars Compacta
SNr: Substantia Nigra Pars Reticulata
tdT: tdTomato (fluorescent protein)
TRAP: Targeted Recombination in Active Populations
WD: Withdrawal

